# fHow ATP and dATP act as molecular switches to regulate enzymatic activity in the prototypic bacterial class Ia ribonucleotide reductase

**DOI:** 10.1101/2021.07.31.454598

**Authors:** Michael A. Funk, Christina M. Zimanyi, Gisele A. Andree, Allison E. Hamilos, Catherine L. Drennan

## Abstract

Class Ia ribonucleotide reductases (RNRs) are subject to allosteric regulation to maintain the appropriate deoxyribonucleotide levels for accurate DNA biosynthesis and repair. RNR activity requires a precise alignment of its α_2_ and β_2_ subunits such that a catalytically-essential radical species is transferred from β_2_ to α_2_. In *E. coli*, when too many deoxyribonucleotides are produced, dATP binding to RNR generates an inactive α_4_β_4_ state in which β_2_ and α_2_ are separated, preventing radical transfer. ATP binding breaks the α−β interface, freeing β_2_ and restoring activity. Here we investigate the molecular basis for allosteric activity regulation in the prototypic *E. coli* class Ia RNR. Through the determination of six crystal structures we are able to establish how dATP binding creates a binding pocket for β on α that traps β_2_ in the inactive α_4_β_4_ state. These structural snapshots also reveal the numerous ATP-induced conformational rearrangements that are responsible for freeing β_2_. We further discover, and validate through binding and mutagenesis studies, a previously unknown nucleotide binding site on the α subunit that is crucial for the ability of ATP to dismantle the inactive α_4_β_4_ state. These findings have implications for the design of allosteric inhibitors for bacterial RNRs.

## Introduction

Ribonucleotide reductases (RNRs) are key enzymes in DNA biosynthesis and repair. They use radical-based chemistry to convert ribonucleotides into deoxyribonucleotides, providing the only *de novo* route for deoxynucleotide production. Due to these functions, RNRs are targets for anti- cancer, anti-viral and antibiotic therapies (Aye et al., 2015; Greene et al., 2020). The best studied RNR is the class Ia RNR from *Escherichia coli. E. coli* class Ia RNR, like the human RNR, requires two dimeric subunits for activity; the α_2_ subunit houses the active site and the allosteric sites, and the β_2_ subunit houses the radical cofactor (**Fig. 1**) (Brown and Reichard, 1969a; Eriksson et al., 1997; Nordlund et al., 1990; Uhlin and Eklund, 1994). On every round of turnover, these subunits come together to form an active α_2_β_2_ complex (Kang et al., 2020), that allows for the transfer of the radical species from β to α, affording ribonucleoside diphosphate reduction (Minnihan et al., 2013).

**Figure 1.**
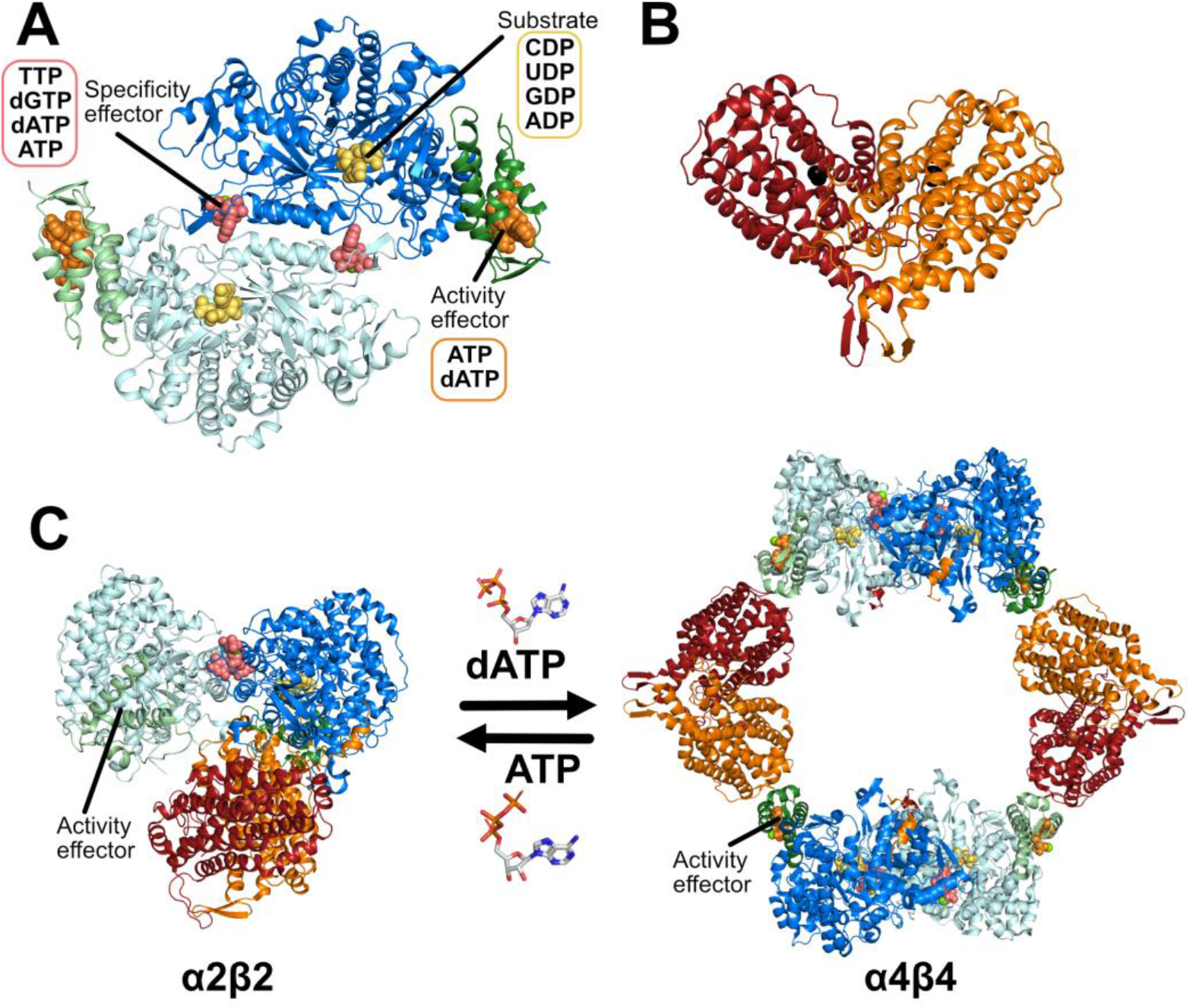
Structures of the *E. coil* class Ia RNR. The individual α and β subunits each form homodimers, shown in cartoon representation. (A) The α_2_ subunit (in cyan and blue, W28A α_2_ structure shown) binds NDP substrates in the active site and ATP and dNTP effectors at the allosteric sites (shown as spheres). The N-terminal regulatory “cone” domain containing the allosteric activity site is colored in green. (B) β_2_ from the structure of the active α_2_β_2_ complex (PDB: 6W4X) (C) A compact α_2_β_2_ structure is required for catalytic activity (PDB: 6W4X). This complex is in equilibrium with the free subunits and an inactive α_4_β_4_ complex (PDB: 5CNS) which is formed when dATP binds to the allosteric activity site. The presence of ATP promotes formation of the active complex.

RNRs act on all four ribonucleoside diphosphates with specificity regulated allosterically by the binding of specificity effectors (ATP/dATP, dGTP, TTP) to the specificity site at the dimer interface of α_2_. In particular, ATP and dATP favor CDP or UDP reduction, TTP binding favors GDP reduction, and dGTP binding favors ADP reduction (Fig. S1) (Brown and Reichard, 1969b; Von Doebeln and Reichard, 1976; Zimanyi et al., 2016). By the use of specificity regulation, one enzyme can provide all four deoxyribonucleotides in the proper ratios required for the fidelity of DNA replication.

Class Ia RNRs are also subject to allosteric activity regulation in which the binding of activity effectors ATP and dATP to the ‘activity site’ in an N-terminal region of the α_2_ subunit, referred to as the cone domain (Aravind et al., 2000) (residues 1-95) (**Fig. 1**), causes the catalytic activity to increase or decrease, respectively. This regulation maintains the appropriate ratios of ribonucleotides to deoxyribonucleotides in the cell (Brown and Reichard, 1969b). Previous work has provided insight into the mechanism of allosteric activity regulation for *E. coli* class Ia RNR, showing that increasing dATP concentrations drives the interconversion of an active α_2_β_2_ state to an inactive state α_4_β_4_ ring-like state through the binding of dATP to the cone domain (**Fig. 1**) (Ando et al., 2011; Rofougaran et al., 2008). This α_4_β_4_ state is inactive because β_2_ subunits are too far (over 60-Å) from α_2_ for radical transfer to occur, effectively turning RNR off (**Fig. 1**) (Ando et al., 2011; Zimanyi et al., 2012). ATP reverses this inhibition by binding to the cone domain and shifting the conformational equilibrium away from α_4_β_4_ toward α_2_β_2_ (Ando et al., 2011). Importantly, RNR variants in which residue substitutions prevent formation of the α_4_β_4_-ring are no longer allosterically regulated by dATP (Chen et al., 2018), indicating that dATP-induced α_4_β_4_-ring formation is causative of the allosteric inhibition. Previous structural studies have indicated that ATP and dATP bind to the same site within the cone domains of α_2_: a structure of *E. coli* class Ia RNR in the α_4_β_4_ state co-crystallized with dATP (Zimanyi et al., 2016, 2012) shows the nucleotide in the same site that was occupied by AMP-PNP (an ATP mimic) in a structure of α_2_ (Eriksson et al., 1997), raising the question of how the binding of nucleotides that differ by one hydroxyl group can lead to such dramatic oligomeric state changes.

Although eukaryotic class Ia RNRs do not form α_4_β_4_-rings, the rules of allosteric activity regulation are the same: dATP turns RNR off and ATP turns it back on (Rofougaran et al., 2006). Structural data show that ATP and dATP bind to the same location in the α_2_ cone domains in human RNR as observed for *E. coli* (Brignole et al., 2018; Fairman et al., 2011), but human RNRs form α_6_- rings (Ando et al., 2016; Brignole et al., 2018; Fairman et al., 2011; Rofougaran et al., 2006). Activity of α_6_ rings in the presence of β_2_ seems to depend of the stability of the α_6_ rings, with dATP-bound α_6_ rings being stable and inactive (Ando et al., 2016; Brignole et al., 2018). Data suggest that β_2_ is prevented from interacting with α_2_ in manner that affords radical transfer when α_2_ is part of a stable ring (**Fig. S2**) (Ando et al., 2016). However, α_6_ rings formed with ATP are unstable and active in the presence of β_2_ (Ando et al., 2016). Thus, for both human and *E. coli* RNR, dATP binding in the cone domain appears to afford ring stability whereas ATP affords ring instability. The molecular basis of this stability/instability is unknown in both cases.

Given the desire to inhibit RNRs as part of anti-cancer or antibiotic therapies, here we provide insight into the natural mechanism of RNR inhibition by dATP, using the well-studied *E. coli* class Ia RNR system. Since subunit interfaces are different in the dATP-inhibited structures of human and bacterial RNRs, these investigations could lead to the development of bacterial-specific RNR inhibitors, a long-standing goal. Toward these ends, we present six crystal structures and accompanying binding and mutagenesis data to investigate the question of how dATP binding stabilizes an approximately 525-Å^2^ α-β interface of the dATP-inhibited α_4_β_4_ ring of *E. coli* class Ia RNR, and how ATP binding dismantles that interface. We find that the molecular mechanism is exquisitely complex and involves a previously unknown second ATP binding site in the cone domain.

## Results

### Overall structures of α_2_ with dATP and ATP are similar to previous α_2_ structure with AMP-PNP

We have obtained a 2.62-Å resolution co-crystal structure of α_2_ with ATP and a 2.55-Å resolution structure with dATP (**Tables S1-S3**). Both structures have nucleotides bound in the specificity and activity sites (**Fig. S3, S4**). Neither structure has substrate. With the exception of the cone domain, there are few overall structural changes between the ATP- and dATP-bound free α_2_ structures with each other or with the previous AMP-PNP structure (**Fig. 2**) (Eriksson et al., 1997). The Cα root-mean-squared deviation (RMSD) for each structure comparison is shown in **Table S4**.

**Figure 2.**
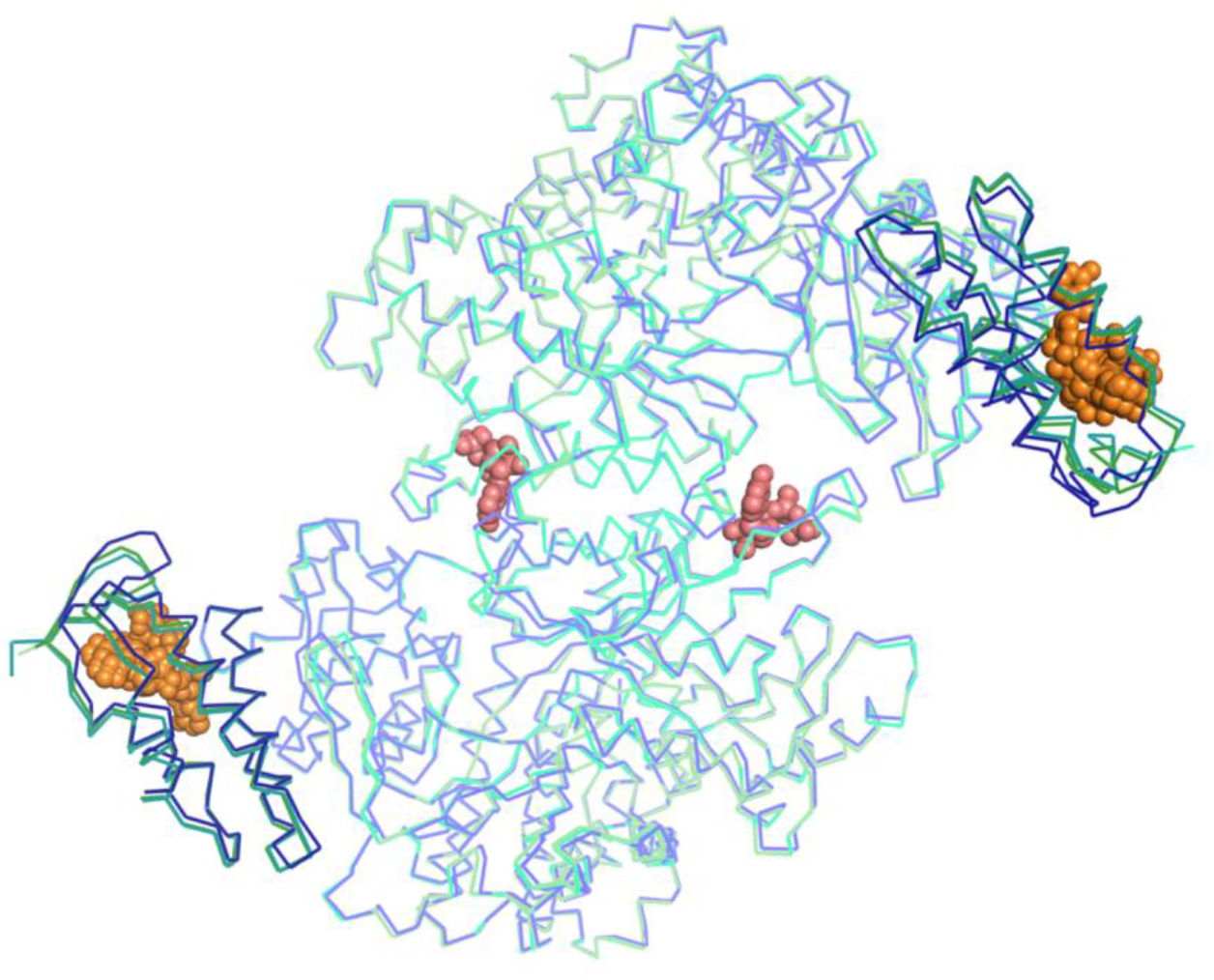
Overall structures of α_2_-dATP and α_2_-ATP agree with that of the previously reported structure of the *E. coli* class Ia RNR α_2_ subunit. The regulatory cone domain is colored in a darker shade of the overall structure. Overlay of α_2_ with AMP-PNP bound at the allosteric activity site (PDB ID: 3R1R) colored in blue/dark blue, α_2_-dATP colored in teal/dark teal (this work) and α_2_-(ATP)_2_ colored in light green/dark green (this work) with nucleotides shown as spheres.

### dATP binds similarly to activity site regardless of the presence of β_2_

The cone domain is at the N-termini of the α_2_ subunits and is comprised of a β-hairpin and a four-helix bundle. We find that dATP binds to the cone domains of the α_2_ structure in an analogous manner as was observed previously in the structure of the dATP-inhibited α_4_β_4_ state (Zimanyi et al., 2016). The α_2_ structure presented here is of higher resolution and contains an intact dATP molecule (the dATP molecule was hydrolyzed to dADP during α_4_β_4_ crystallization), and thus affords a more complete picture of dATP binding to the *E. coli* class Ia RNR enzyme **(Fig. 3A,B, Fig. S4A,B**). Specificity for the adenine base of dATP is generated by hydrogen bonds between the side chain carboxylate of E15 (acceptor) and N6 (donor) and the backbone amide NH of N18 (donor) and N1 (acceptor). The base is additionally held in place by packing interactions with residues V7 and I17 (β-hairpin), I22 (helix 1), and F49 and I58 (helix 3) (**Fig. 3D**). The deoxyribose moiety sits between helix 1 and 3 of the four-helix bundle with a single hydrogen bond made between O3′ and helix 3 residue H59. β-hairpin residues K9 and R10 and helix 4 residue K91 provide charge neutralization and electrostatic interactions with the phosphates of dATP, which also coordinate a Mg^2+^ ion. The only difference in the coordination environment between the α_2_ and α_4_β_4_ structures is the side chain of T55, which adopts different rotamer conformations that form different hydrogen bonds (**Fig. 3A,B**). This variation may be due to subtle changes in coordination caused by the loss of the gamma phosphate of dATP over the time course of the α_4_β_4_ crystallization.

**Figure 3.**
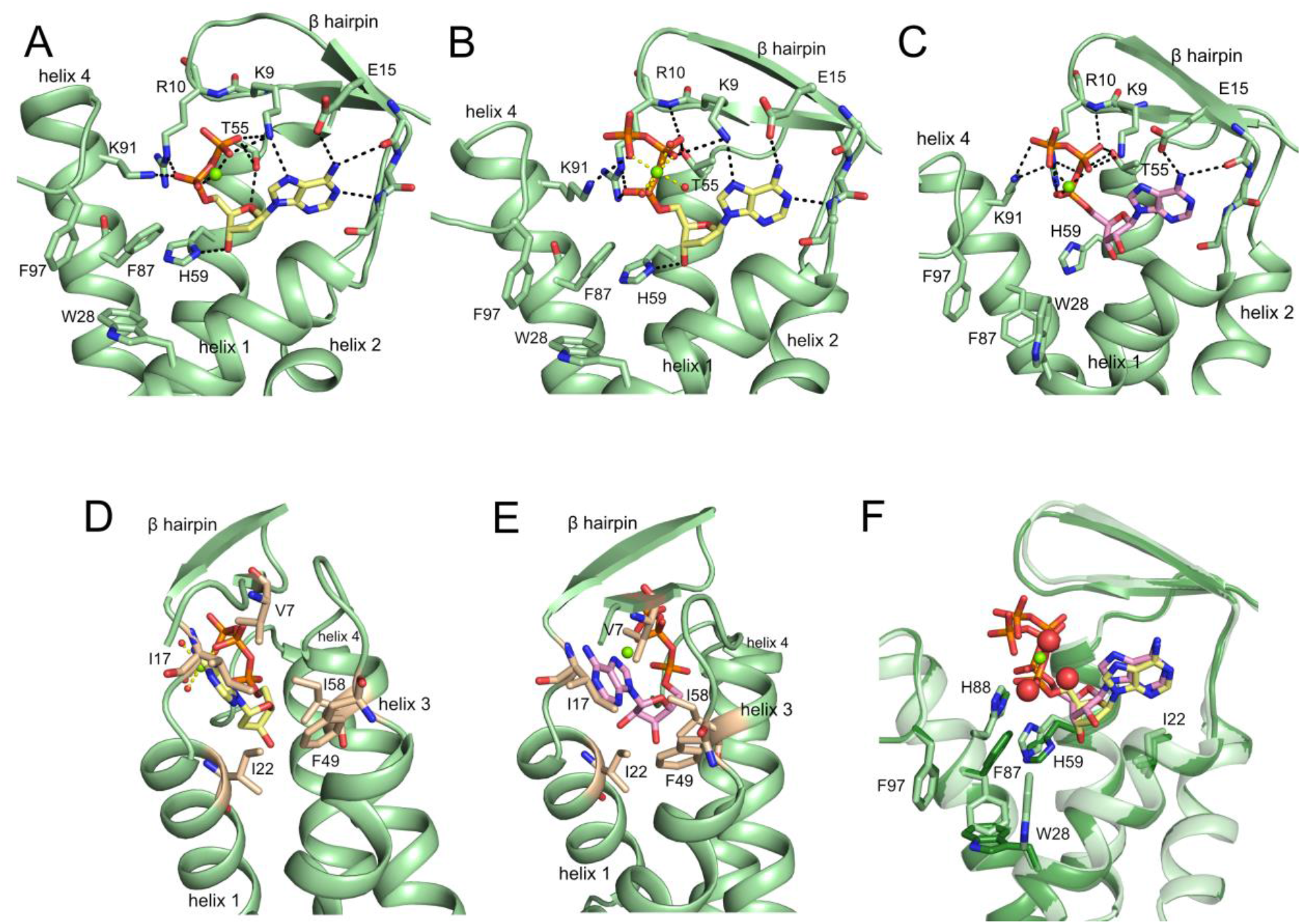
Binding of dATP and ATP to the previously identified allosteric activity site in the cone domain in various *E. coli* class Ia RNR structures. (A) dATP-inactivated α_4_β_4_ state of *E. coli* class Ia RNR with a hydrolyzed dATP activity effector bound to the cone domain (PDB: 5CNS). Structure was obtained in the presence of dATP but dATP hydrolyzed to dADP (yellow) over the time course of the crystallization. (B) α_2_ with dATP bound (yellow) to the activity site in the cone domain. (C) α_2_-(ATP)_2_ with ATP (pink) bound to site 1 (the canonical site) in the activity site in the cone domain. (D) Hydrophobic contacts (residues in tan) for dATP (yellow) in α_2_-dATP structure. (E) Hydrophobic contacts (residues in tan) for ATP (carbons in pink) in α_2_-dATP structure. (F) Superposition of the cone domains of α_2_-(ATP)_2_ (light green, one ATP in pink shown, occupying site 1) and α_2_-dATP (dark green, dATP in yellow). Water molecules are shown as red spheres, Mg^+2^ ions in green. Residues that contact nucleotides or move upon nucleotide binding are labeled.

### There are two distinct ATP binding sites in the cone domain of α_2_

Surprisingly, we find two molecules of ATP bound to the cone domain **(Fig. 4A, S4C,D**). The first binding site for ATP is the same activity site to which dATP binds (we will call this site 1). The second site (site 2) is generated through a series of conformational changes afforded by the binding of the first ATP molecule. This second nucleotide-binding site does not exist in the apo form of α_2_ nor in dATP-bound α_2_.

**Figure 4.**
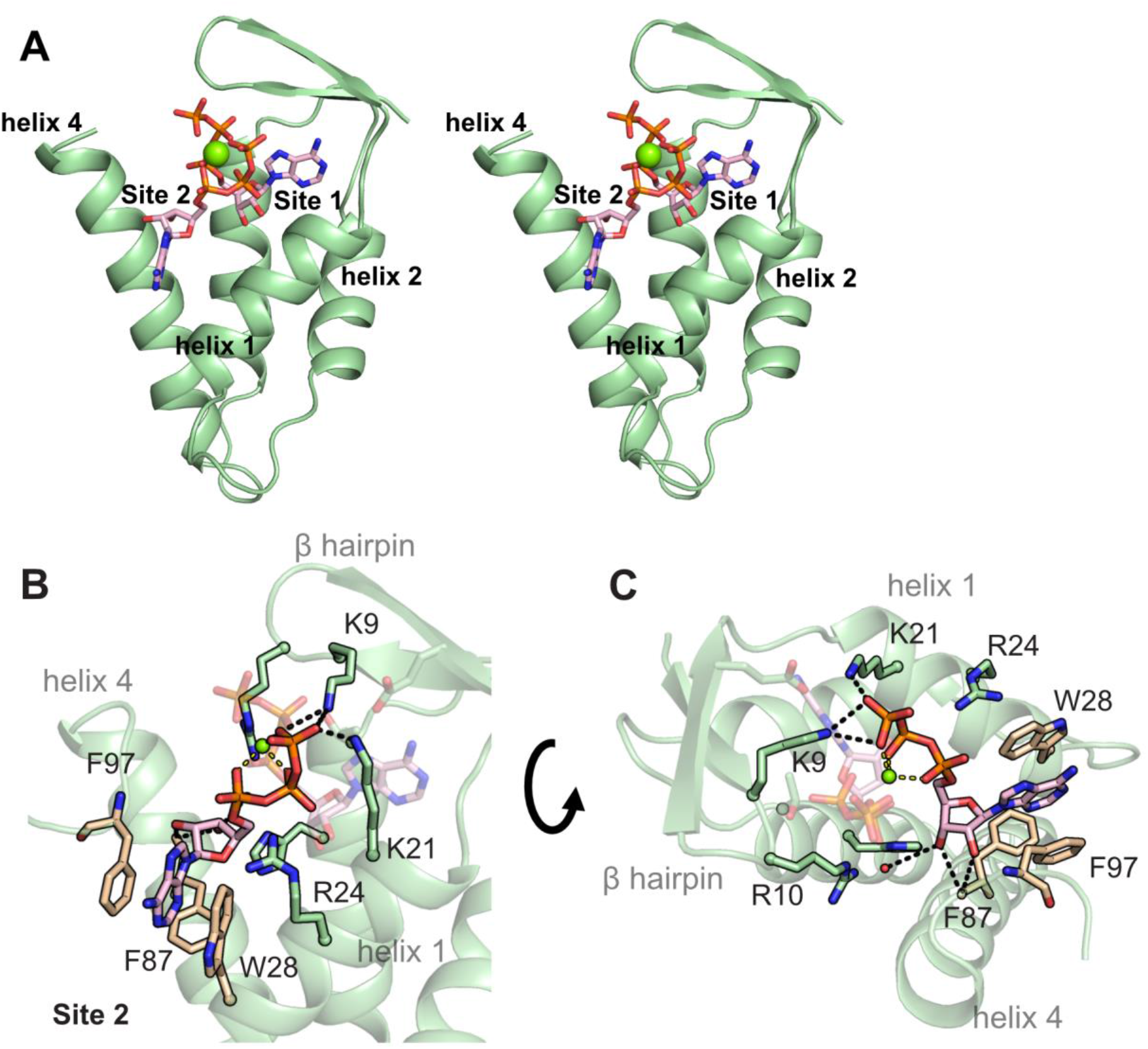
A second binding site for ATP is discovered in the cone domain of the α_2_ subunits of *E. coli* class Ia RNR. (A) Stereoview of the cone domain with two molecules of ATP (α_2_-(ATP)_2_) in site 1 and site 2. (B) Interactions of ATP in site 2 of the α_2_-(ATP)_2_ structure. (C) Interactions of ATP in site 2 of the α_2_-(ATP)_2_ structure, showing site specificity for ribonucleotides. Hydrogen bonds are shown as black dashes, and Mg^2+^ contacts to phosphate groups are shown as yellow dashes. Hydrophobic residues in tan.

The first ATP molecule binds to site 1 by making very similar interactions as dATP (**Fig. 3**). The hydrogen bonds to the adenine base are identical (**Fig. 3B,C**). The presence of the 2′-hydroxyl group results in a slight adjustment upward of the ribose ring of ATP compared to the ribose in dATP due to the packing of the ribose against I22 of helix 3 (**Fig. 3D, E**). This subtle shift of the ribose results in the loss of a hydrogen bond between the ribose 3′-hydroxyl group and H59, whose side chain flips 90° and now hydrogen bonds to a water molecule (**Fig. 3F**). The phosphates of ATP also sit slightly higher in the activity site, with favorable hydrogen-bonding/electrostatic interactions being made by K9, R10, and K91 (**Fig. 3C**). As a result of these subtle movements, the β-hairpin is slightly shifted (**Fig. 3F**).

The second nucleotide-binding site (site 2) is directly adjacent to site 1 within the cone domain, sandwiched between helix 1 and helix 4 (**Fig. 4A**). The phosphate groups of the two ATP molecules create an octahedral coordination environment around a central Mg^2+^ ion. The additional negative charge from the second triphosphate moiety is stabilized by electrostatic/hydrogen-bonding interactions from K9 and K21 (**Fig. 4B**). R24 is also within 4 Å of the phosphate groups, although it does not make direct contacts in the crystal structure. The site 2 ATP ribose makes hydrogen-bonding interactions to the backbone carbonyl of F87 through O2′ and O3′, suggesting specificity at this site for ribonucleotides (**Fig. 4C**). Unlike in site 1, there are no specific contacts to the adenine base; instead, it is held in a hydrophobic pocket between helix 1 and 4.

### Creation of second ATP binding site involves a coordinated movement of three side chains

The transition from the apo- or dATP- to the two ATP-bound form of the cone domain requires a dramatic and coordinated shift of three key residues—H59, F87 and W28—all three of which adopt different rotamer conformations upon ATP binding (**Fig. 5, Fig. S5**). As mentioned above, the side chain of H59 hydrogen bonds to O3′ of dATP but is flipped out of the activity site when ATP is bound. This new position of the H59 side chain is in close proximity to the side chain of F87, and F87 must adopt a flipped-down position. This flipped-down position of F87 brings its side chain into the pocket occupied by the W28 side chain, causing W28 to flip 90° sideways. The net result of these side chain movements is formation of a new nucleotide-binding site in the cone domain. F87 movement creates the binding pocket for the ribose of the second ATP molecule (**Fig. 5, Fig. S5**) and W28 movement creates a cavity for the base of the second ATP. The adenine base is nicely sandwiched between side chains of W28 and F97 as a result of W28’s movement.

**Figure 5.**
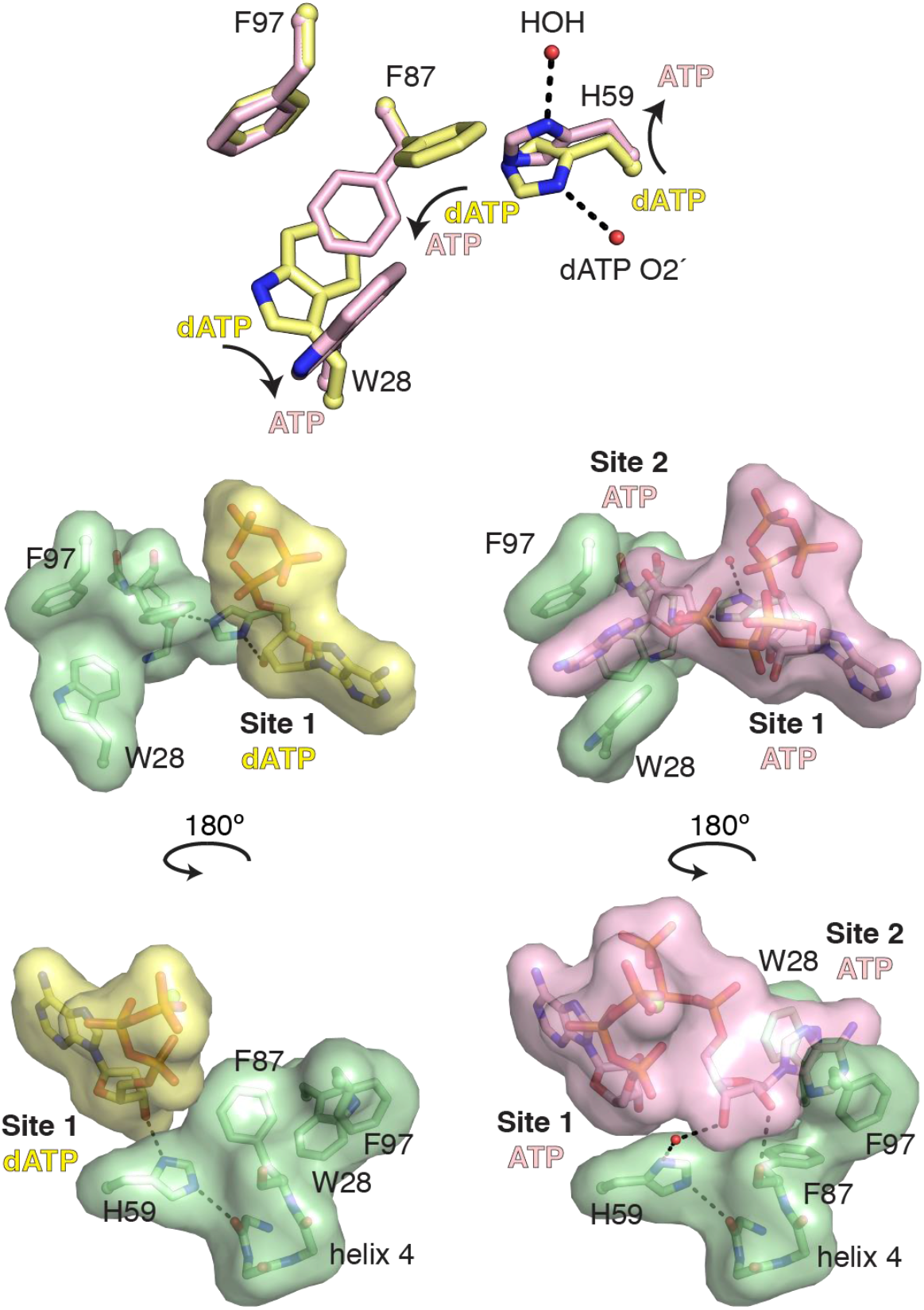
A conformational switch involving His59 is required to generate the second ATP site in the cone domain. (A) W28, H59, and F87 undergo rotamer shifts upon the binding of two ATP molecules. F97 does not change, as dATP only binds in site 1. (B & C) Space filling models of the interaction of the dATP/ATP molecules with helix 4 and the W28, H59, F87, and F97 residues.

### Equilibrium binding assays are consistent with two higher affinity ATP binding sites at the activity site and one lower affinity ATP binding site at the specificity site

The presence of two ATP molecules at the activity site was unexpected. Previous ultrafiltration binding studies by Ormö and Sjöberg (Ormö and Sjöberg, 1990) indicated that two ATP molecules bind per α, which was interpreted as signifying the binding of one ATP molecule to the specificity site and one to the activity site. The α_2_-ATP structure shows three ATP molecules per α in total: one ATP in the specificity site and, as described above, two ATP molecules in the activity site. To pursue the possibility that a lower-affinity ATP binding site might have gone undetected if the ATP concentration was not high enough in the previous ultrafiltration binding studies, we have re-run these assays using a higher concentration of ^3^H-ATP and using non-linear regression to analyze the data. In this assay, the concentration of free and bound ^3^H-ATP is determined after separation of the protein by centrifugation in a spin filter, allowing for determination of equilibrium binding parameters. The resulting binding curves from multiple experimental runs were fit with a one-state binding model that assumes all binding sites are equivalent. This simplifying assumption is necessary due to the impracticality of fully sampling the binding curve. Using this approximation, the *K*_d_ for ATP binding at 25 °C was estimated to be 158 ± 37 μM with the maximum number of binding sites being 6.8 ± 0.6 per α_2_ (**Fig. 6**). This number is consistent with this structure, which shows 3 ATP molecules per α (6 ATP molecules per α_2_).

**Figure 6.**
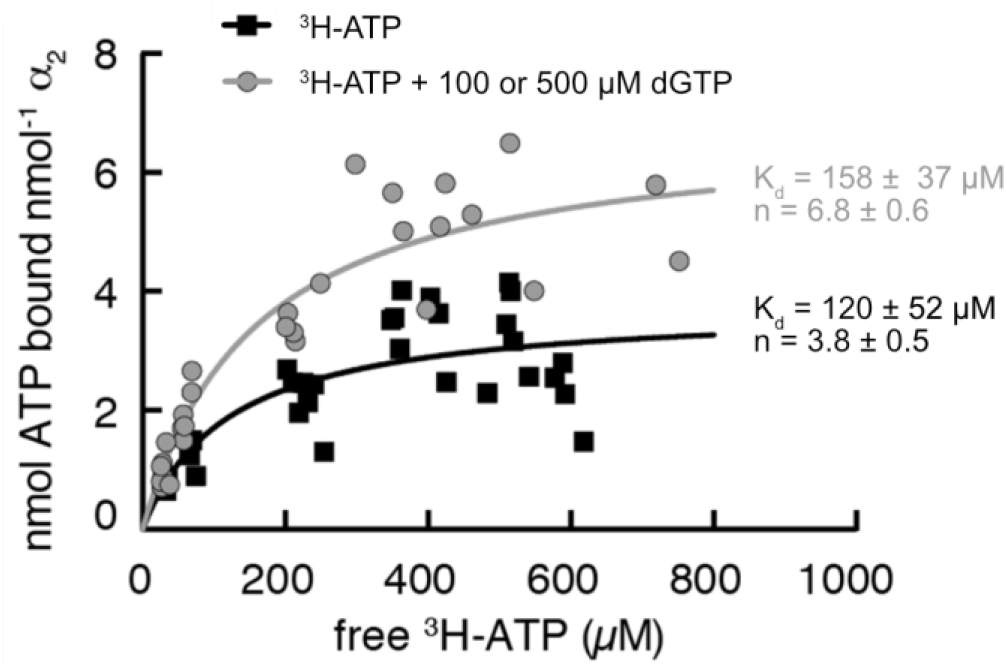
ATP binding curves measured by the ultrafiltration method indicate the binding of multiple ATP molecules per α_2_ subunit of *E. coli* class Ia RNR. The ultrafiltration method described by Ormö and Sjöberg (Ormö and Sjöberg, 1990) was used with modifications to determine the equilibrium binding parameters for ATP. Each point is an independent measurement. The sample consisted of 7-20 μM α_2_, ^3^H- ATP binding to α_2_ alone is in gray circles. ^3^H-ATP binding to α_2_ in the presence of 100 or 500 μM dGTP is in black squares. These dGTP concentrations saturate the specificity site allowing for the determination of activity-site-only binding parameters. The amount of bound nucleotide was found by subtracting the amount of free nucleotide from total nucleotide and resulting binding curves from multiple experimental runs were fit with a one-state binding model that assumes all binding sites are equivalent. Using this approximation, the *K*_d_ for ATP binding at 25 °C was estimated to be 158 ± 37 μM with the maximum number of binding sites being 6.8 ± 0.6. In the presence of dGTP, the *K*_d_ for ATP at the allosteric activity site was measured to be 120 ± 52 μM with a maximum of 3.8 ± 0.5 binding sites. These data are consistent with the binding of two ATP molecules at each activity site.

To verify the presence of multiple binding sites for ATP at the activity site, the ultrafiltration experiment was repeated in the presence of 100 or 500 μM dGTP, which should bind at full occupancy to the specificity site given that the *K*_d_ for dGTP at the allosteric specificity site is 0.77 μM (Ormö and Sjöberg, 1990). In the presence of dGTP, the *K*_d_ for ATP was measured to be 120 ± 52 μM with a maximum of 3.8 ± 0.5 binding sites per α_2_ (**Fig. 6**). These data are consistent with the binding of two ATP molecules in each cone domain. No difference in binding was observed at the two different dGTP concentrations, suggesting that the specificity site is indeed saturated with dGTP under these conditions. Importantly, these data also indicate that the binding sites in the cone domain are the higher affinity sites for ATP, further suggesting that the initial binding study was most likely reporting on two ATP molecules binding to the activity site. We can now also explain the previous observation that ATP binding is cooperative (Brown and Reichard, 1969b; Ormö and Sjöberg, 1990), which was hard to explain if ATP was binding to sites that are more than 40 Å apart, but easy to explain if ATP molecules are coordinated by a single Mg^2+^ ion and if the binding of the first ATP creates the binding site for the second ATP, as it appears to do. Thus, in terms of the question of how the difference of one hydroxyl group (ATP vs dATP) can destabilize the α_4_β_4_ ring and shift the equilibrium back to an active α_2_β_2_ state, these structures and these binding data suggest that it is the one extra hydroxyl group of ATP, and one whole extra molecule of ATP, which are responsible for α_4_β_4_ destabilization.

### Identity of nucleotide in site 1 can be uncoupled from nucleotide binding in site 2 through a W28A substitution

With the confirmation from the binding assays that two ATP molecules bind to the activity site, we next sought to test the importance of ATP binding at site 2 to the overall activity regulation of *E. coli* class Ia RNR. We generated individual alanine variants of residues W28, F87, and F97, which comprise the second ATP binding site (**Fig. 4**). As described above, F87 and W28 side chains move between rotamer conformations that alternatively block the second ATP from binding (residue positions in yellow in **Fig. 5**) and support the second ATP binding (residue positions in pink in **Fig. 5**), as signaled by the position of H59, which moves in response to the presence of ATP versus dATP in site 1. We reasoned that substitutions of either F87 or W28 with Ala would create room for ATP binding in site 2 regardless of the position of H59 and thus be independent of the presence of ATP in site 1. In other words, F87A and W28A substitutions would be expected to uncouple the binding of ATP to site 2 from the identity of the nucleotide effector in site 1. To test this idea, we obtained structural data for W28A-α_2_ with ATP, with dATP/ATP, and with dATP/GTP (**Table S1, S3**). First, we wanted to determine if ATP could enter site 2 with dATP in site 1 in a W28 construct, so we grew crystals in the presence of a mixture of 3 mM to 5 mM ATP and 1 mM dATP. The resulting 3.40-Å-resolution structure revealed clear density for two nucleotides in the activity site, but the identity of the site 1 nucleotide was ambiguous given the low resolution of the structure (**Fig. S6A,B**). Due to the lack of clarity about the identity of the nucleotide in site 1, this structure could not be used as evidence that a W28A substitution leads to site 1, 2 uncoupling. Unfortunately, attempts to obtain a high-resolution structure with ATP and dATP have so far been unsuccessful.

Thus, we tried a different approach, co-crystallizing W28A α_2_ with 1 mM dATP and then soaking with 3 mM GTP. As described above, site 1 is specific for a nucleotide with an adenine base (see **Fig. 3**) and cannot accept GTP. However, site 2 shows no interactions with adenine base of ATP except for hydrophobic packing and thus should be able to accommodate other ribonucleotides (see **Fig. 4**). Excitingly, the resulting structure, albeit at the modest resolution of 3.55-Å-resolution, does show density in both site 1 and site 2 (**Fig. S6C,D Table S1, S3)**. With the W28A substitution, the side chain of F87 is positioned down, creating the site 2 binding pocket, and allowing for nucleotide binding, despite the fact that H59 has not moved (**Fig. 7**). The density in site 2 is consistent with GTP binding in a very similar conformation as found for ATP in the WT α_2_-(ATP)_2_ structure (**Fig. S6D,H**). We also confirmed that W28A substitution does not alter ATP binding in site 2 by determining a crystal structure of W28A α_2_ with ATP as the only effector at 2.6-Å resolution (**Table S1, S3**). We find that the W28A substitution results in a more open pocket for ATP binding at site 2, but no structural perturbations in the bound ATP molecule are apparent (**Fig. S6E,F,G,H**). All site 1 residues and site 2 residues F87 and F97 are identical to WT α_2_-(ATP)_2_ (**Fig. 7, S6E,F,G,H**). Therefore, the W28A substitution appears successful in uncoupling nucleotide binding in site 2 from the identity of the nucleotide in site 1 without substantially altering ATP binding to site 2.

**Figure 7.**
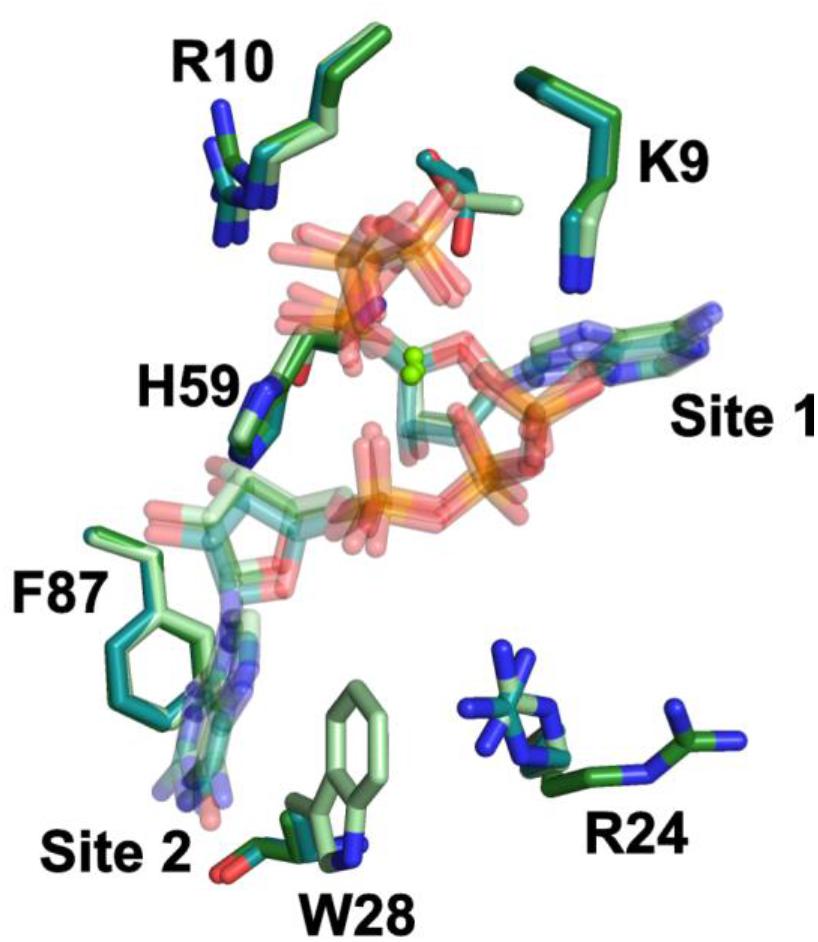
Overlay of residues involved in nucleotide binding in both site 1 and 2. The residues and bound nucleotides of wild-type α_2_-(ATP)_2_ are colored in light green, W28A α_2_-(ATP)_2_ are colored in dark green, and W28A α_2_-(dATP/GTP) are in teal. Nucleotides are shown with transparency, residues and sites are labeled.

Although not the focus of this report, the W28A α_2_-(ATP)_2_/CDP structure at 2.6-Å resolution structure also provides a higher-resolution picture of the active site bound to the CDP/ATP substrate/effector pair (**Fig. S7**). The hydrogen bonding pattern that affords substrate specificity is the same as that described previously (Zimanyi et al., 2016).

### Substitutions W28A, F87A, and F97A at the second ATP binding site disrupt activity regulation

With the knowledge that the W28A substitution uncouples nucleotide identity at site 1 from that of site 2, we proceeded to investigate the relevance of site 2 to allosteric regulation of activity. If site 2 is not important in either dATP-induced activity down-regulation or ATP-induced activity up-regulation, then the W28A RNR variant should behave like WT protein. As controls, we also investigated an F87A RNR variant, which should also uncouple site 1 and site 2, and an F97A RNR variant that should not lead to site 1, 2 uncoupling, but would be expected to decrease the affinity of ATP binding in site 2 due to loss of the stacking interaction of the adenine base with the F97 side chain (**Fig. 4**). Our results described below indicate that site 2 is, in fact, critical for the ability of RNR to be regulated by the ratio of dATP to ATP, but is not important for activity under ATP-only conditions, and is not important for the ability of dATP to down-regulate RNR when no ATP is present (**Fig. 8**). In particular, our data show that under activating conditions (3 mM ATP as specificity and activity effector), these three enzyme variants had activities comparable to wild-type α_2_ with CDP as substrate (**Fig. 8A**). In the presence of inhibitor dATP, F97A was inactivated to a degree similar to wild-type α_2_ (5-10% of maximal), and the activity of W28A and F87A was reduced somewhat less (20-25% of maximal) (**Fig. 8A**). Overall activity regulation in these three mutants is thus only moderately perturbed when ATP and dATP are used as effectors in isolation.

**Figure 8.**
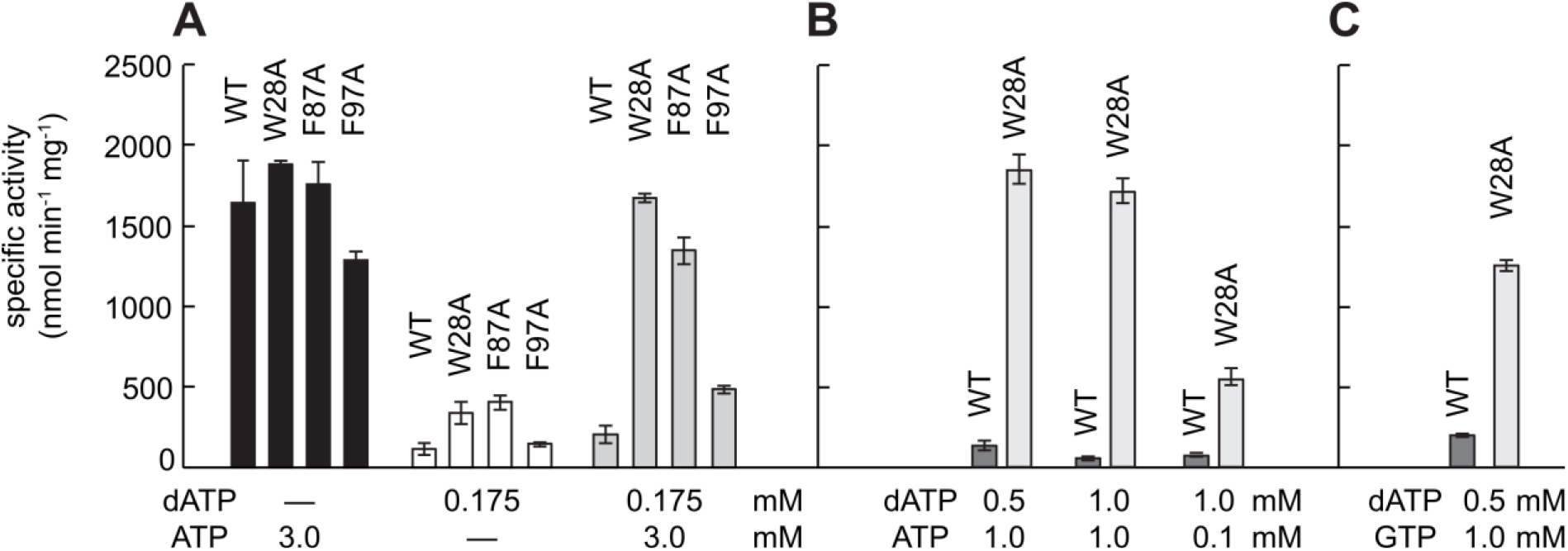
Substitutions at the second ATP binding site (W28A, F87A, and F97A) disrupt allosteric activity regulation. (A) Specific activity of RNR assessed for different ratios of allosteric activity effectors ATP and dATP for WT and three RNR variants. Substitution of W28, F87, and F97 with alanine leads to RNR variants that are similarly active with 3.0 mM ATP (black bars) as WT, deactivate with 0.175 mM dATP like WT (white bars), although not as fully as WT, and unlike WT, re-activate when 3.0 mM ATP is added to 0.175 mM dATP (grey bars). (B) Specific activity of WT and the W28A variant RNR assessed for a wider range of ATP and dATP conditions. Whereas WT RNR (dark grey bars) is largely inactive when the dATP concentration is 0.175 mM or higher, regardless of the amount of activator ATP, the W28A variant (light gray bars) is very sensitive to the addition of ATP and shows activity even when ATP concentrations are ten-fold less than dATP concentrations. (C) Specific activity of WT and the W28A variant RNR in the presence of inhibitor dATP (0.5 mM) with activator ATP replaced by 1.0 mM GTP. The substitution of W28 for alanine appears to alter the specificity of site 2 for ATP such that GTP can now upregulate enzyme activity. dATP and ATP concentrations are shown below the graph. Error bars shown are standard deviations.

To evaluate the activating effect of ATP in combination with dATP, we tested a condition containing 3 mM ATP and 0.175 mM dATP (**Fig. 8A**), which are physiologically relevant concentrations for *E. coli.* With 3 mM ATP and 0.175 mM dATP, wild-type α_2_ and F97A activity are stimulated only slightly relative to 0.175 mM dATP alone, and remain low relative to 3 mM ATP alone. In contrast, W28A and F87A variants in 3 mM ATP and 0.175 mM dATP both recover near maximal activity under these conditions. This finding is consistent with the ability of ATP to bind to site 2 in W28A and F87A variants even when dATP is bound in site 1, and is consistent with the importance of ATP binding in site 2 to the up-regulation of RNR activity. Notably, our structural data suggest that site 2 is restricted to ribonucleotides and these data support this conclusion. If dATP was binding to site 2 in either the W28A or F87A RNR variant, then these proteins would not show these WT-like levels of dATP inactivation. Therefore, site 2 does not appear to be important to the ability of RNR to be turned off by dATP, but is important for the ability of RNR to be turned on by ATP.

We further tested the W28A variant under different ratios and concentrations of dATP and ATP (**Fig. 8B**) and found that the W28A variant is fully active in the presence of 1.0 mM ATP under extremely high concentration of dATP (0.5 mM or 1 mM). With a K_d_ of ∼ 6 μM for dATP binding (Brown and Reichard, 1969b; Ormö and Sjöberg, 1990) and ∼120 μM for ATP binding to site 1, concentrations of 1.0 mM dATP and 1.0 mM ATP should be completely inhibitory, but W28A RNR is fully active. Activity does start to go down when the concentration of ATP is lowered to 0.1 mM and dATP is kept at 1.0 mM. Again, these data indicate the importance of site 2 to successful activity regulation by dATP and ATP.

Given the structural observation that GTP binds to site 2 in W28A-α_2_, we investigated whether GTP can reverse dATP-inactivation, and it can (**Fig. 8C**). Under conditions where WT RNR is inactive (0.5 mM dATP and 1.0 mM GTP), W28A-α_2_ is active, although not as active as W28A-α_2_ with 0.5 mM dATP and 1.0 mM ATP. These results indicate that GTP binding to site 2 can reverse dATP-induced inhibition, but that GTP does not bind as well as ATP to this site. It is very interesting that the presence of a ribonucleotide in site 2, regardless of identity, can restore RNR activity. Of course, under physiological conditions, the concentration of ATP is much higher than the concentrations of other ribonucleoside triphosphates (Bochner and Ames, 1982; Buckstein et al., 2008), such that binding of other NTPs in site 2 is unlikely to be relevant *in vivo*. This result does, however, serve to confirm the importance of site 2 for up-regulation of RNR activity. Again, these data suggest that allosteric regulation by dATP and ATP is not just due to the difference of one hydroxyl group. The hydroxyl group is extremely important, but it is not the whole story. Both the canonical activity site (site 1) and the newly discovered site 2 are critical for allosteric regulation.

### ATP binding leads to an increase in the helix 2 helicity, which in turn destabilizes the α_4_β_4_ structure

We next considered the molecular basis for α_4_β_4_ destabilization by the binding of two molecules of ATP by comparing α_4_β_4_ and α_2_-(ATP)_2_ structures. There are three changes in the cone domain of note when comparing these structures. First, in the presence of the two molecules of ATP, the ^9^KRDG^13^ motif of the β-hairpin of the cone domain is shifted upward ∼2.3 Å (∼6° to ∼7°) at the tip relative to the pivot point at V7 between these two structures (**Fig. 9A**). As mentioned above, as compared to dATP bound in site 1, the ribose of the site 1 ATP and the ATP phosphates sit higher in the activity site, and the site 1 ATP phosphates now share a coordinating Mg^2+^ with the phosphates of the site 2 ATP. The different positioning of phosphate groups alters the position of K9 and R10 of the ^9^KRDG^13^ motif. R10 and K9 both shift up and K9 no longer contacts the adenine base of the nucleotide in site 1 (**Fig. 3A,B**), instead contacting a γ-phosphate of the site 2 ATP (**Fig. 4B,C**). As a result, the ^9^KRDG^13^ motif of the β-hairpin of the cone domain shifts upward. This upward shift results in the second change of note: an approximately equal and opposite downward shift (2.5 Å) of residues Y50-I53 near the base of the β-hairpin (**Fig. 9B**). This downshift of residues Y50-I53, in turn, releases a strain on the residues at the end of helix 2, which can now fold up into an additional helical turn, which is the third change (**Fig. 9B**). Consequently, helix 2 has an extra turn in the α_2_-(ATP)_2_ structure than it does in the dATP-inactivated α_4_β_4_ structure (**Fig. 9C**).

**Figure 9.**
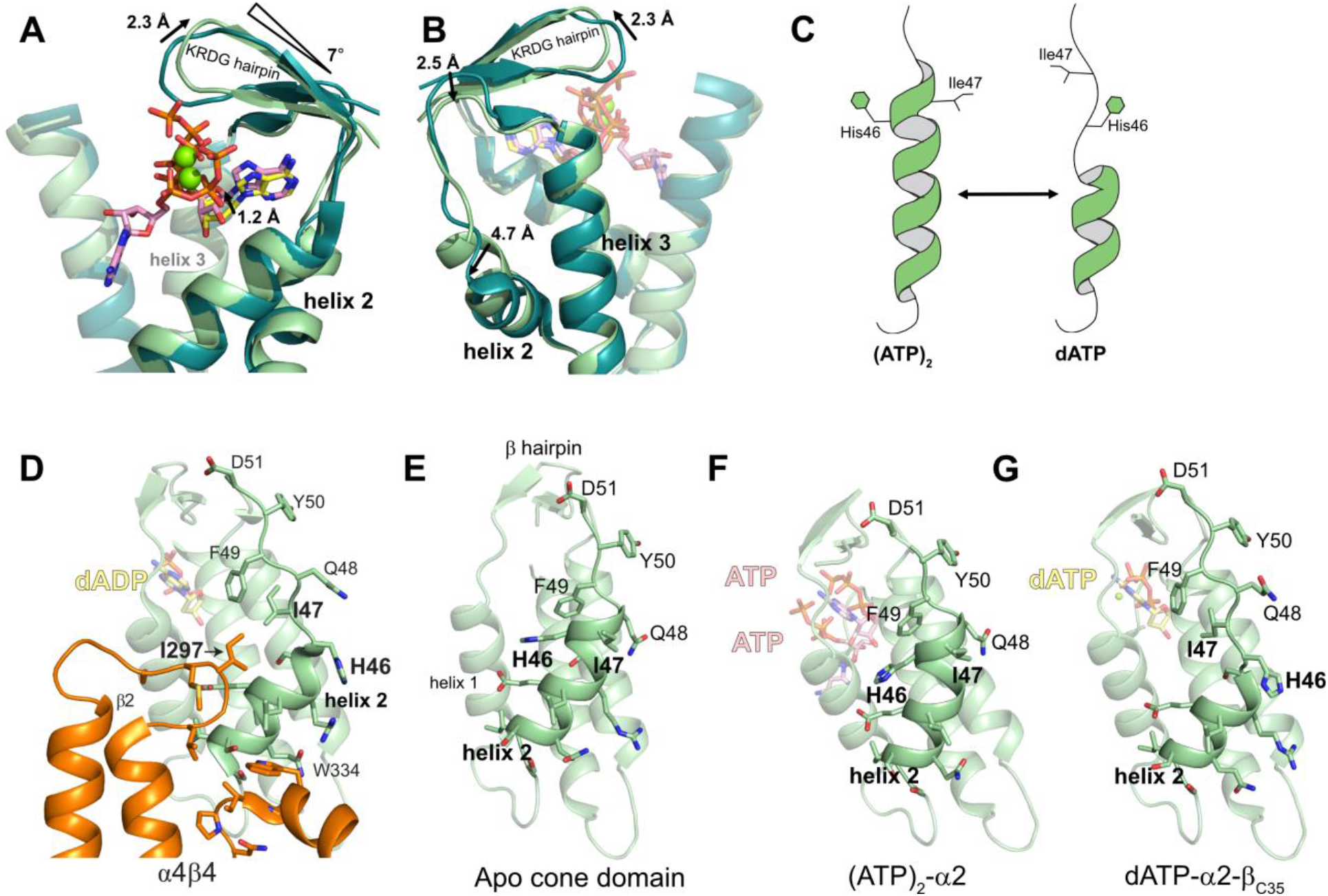
A series of conformational changes induced by the differential binding of dATP versus two ATP molecules results in the creation and destruction of the β-subunit binding surface on α. (A) Differences between dADP-inhibited α_4_β_4_ structure (dark teal, PDB: 5CNS) relative to ATP bound α_2_ structure (light green) are noted in terms of average distance in Å, and the hinge motion in the KRDG hairpin is noted in degrees. Residues Y50-I53 are at the base of the hairpin (not shown). (B) Back view of structural alignment shown in Panel A. Direction of arrows indicates the movement when ATP replaces dATP. (C) Cartoon showing helix 2 conformation with 2 ATP molecules bound compared to the helix 2 conformation with dATP bound. (D) α_4_β_4_ (PDB: 5CNS) showing the interface generated by the unwinding of helix 2. (E) Structure of the apo regulatory cone domain (PDB: 1R1R). (F) α_2_-(ATP)_2_ with helix 2 fully pulled up. (G) dATP-bound α_2_ in the α_2_-βC35 crystal form that allows cone domain movement with helix 2 relaxed.

Thinking about these conformational changes in the opposite direction, the order of events would be: dATP replaces ATP in site 1 and shifts the β-hairpin down; residues Y50-I53 shift up in response; and helix 2 unwinds due to the strain. Thus, the β-hairpin appears to act as a lever responding to the presence of dATP or 2 ATP molecules, alternatively pulling on and unwinding helix 2, or relaxing and refolding helix 2 (**Fig. 9C**). This change in helicity also alters the positions of residues H46 and I47 dramatically.

### dATP binding unwinds helix 2, which creates a binding pocket for residues of the β-subunit, stabilizing an α_4_β_4_ structure

To understand the significance of helix 2 unwinding in the present of dATP, we evaluated the α−β interface in the previously determined α_4_β_4_ structure (Zimanyi et al., 2016, 2012). We find that the unwinding of helix 2 creates a hydrophobic binding pocket for residue I297 of the β subunit to bind, in which I297 packs against I47 (**Fig. 9D**). When helix 2 is fully wound, as it is in the apo structure (**Fig. 9E**, PDB: 1R1R (Eriksson et al., 1997)) and as it is in the α_2_-(ATP)_2_ structure (**Fig. 9F**), there is no binding pocket for I297 of β between helix 1 and 2. The turn of the helix 2 physically blocks the β-binding site. Additionally, polar residue H46 is pointing toward the binding site for I297, making for an unfavorable interaction, and I47 is unavailable to make a favorable one as it is pointing is the opposite direction (**Fig. 9D-F**). Thus, the unwinding of helix 2 of the cone domain of α appears key for the formation of a binding site for the β subunit, creating not only the room for β to bind, but also swapping out an unfavorable interaction (H46) for a favorable one (I47).

Considering the conformational changes in reverse: the binding of two molecules of ATP would destabilize the α_4_β_4_ inactive state by shifting the β-hairpin lever up, releasing the strain on helix 2. As helix 2 refolds, the hydrophobic pocket for residue I297 of β is lost, and the favorable interaction with I47 is replaced with an unfavorable one (H46) to ensure β’s departure. The overall interface is small, ∼525 Å^2^, making it relatively easy for small changes to break α and β apart.

Although the above mechanism of helix 2 unwinding and re-winding in the presence of dATP and ATP, respectively, beautifully explains how the binding surface for the β subunit can alternately be exposed and tucked away, and all ATP-bound structures show helix 2 fully wound (**Fig. S8**), we were puzzled as to why our structure of α_2_-dATP, described above, did not show an unwound helix 2. The structure of dATP-bound α_4_β_4_ shows the unwound helix 2, but we would expect helix 2 unwinding to proceed β binding, since helix unwinding appears to create the β binding site. The lack of helix unwinding in the α_2_-dATP structure cannot be attributed to differential contacts made by dATP, since as noted above, the same contacts are observed in the α_2_ structure as in the α_4_β_4_ structure (**Fig. 3A, B**). Thus, we investigated whether lattice contacts in the crystal might be restricting movement of the β−hairpin in the α_2_-dATP structure, such that helicity of helix 2 would be unchanged, and in fact, residues at the base of the β−hairpin (Y50-I53) are involved in lattice contacts (**Fig. S8)**.

We therefore sought a different crystal form for α_2_ in which the β-hairpin region of the cone domain is less restricted by lattice contacts. Finding new crystal forms for α_2_ has historically been problematic due to the fact that the α_2_ subunits are not very soluble in the absence of the β-subunits. In fact, the first crystal form of α_2_ was obtained through co-crystallization of the α_2_ subunit with a short peptide (residues Y356-L375) that contained the sequence of the β subunit C-terminus (Uhlin and Eklund, 1994). With this in mind, we were able to obtain a new crystal form of α_2_ by fusing the 35 C-terminal residues of β (342-376) to the C terminus of an 8-residue truncated α (α_2_-β_C35_) (**Fig. S9**). The overall structure of this crystal form with dATP bound at 2.10-Å resolution (**Table S1, S2**) is essentially identical to that observed above for free α_2_; however, the cone domain is slightly less restricted by lattice contacts near the β-hairpin (**Fig. S8E**). With the restraints of the crystal lattice relaxed, helix 2 adopts the unwound conformation as in the α_4_β_4_ complex (**Fig. 9G**). The hydrophobic pocket vacated by H46 is left unfilled in this structure, and residues 19-27 (helix 1) and 35-43 (helix 2) that interact with β_2_ in the α_4_β_4_ complex are exposed to solvent. This structure thus demonstrates that specific contacts with β_2_ are not required for helix 2 to be unwound and connects dATP binding in site 1 with helix 2 unwinding.

## Discussion

Allosteric activity regulation in enzymes often involves the movement of side chains in an active site in response to the binding of an allosteric effector in order to either increase or decrease enzyme activity. With this in mind, the incredible oligomeric state change associated with *E. coli* class Ia RNR activity regulation is even more impressive (**Fig. 1C**). *E. coli* class Ia RNR’s mechanism for allosteric regulation of activity can be described as a game of β_2_ keep away, preventing radical transfer and thus activity by keeping β_2_ at arms-length, trapped in a ring structure. But, how is it that the dATP binding stabilizes the trapped ring state whereas, ATP, a molecule that differs only by a single hydroxyl group, allows the ring to fall apart, freeing β_2_. The work presented here suggests that the difference is one hydroxyl group and a whole second molecule of ATP and indicates that protein conformational changes that are involved in controlling ring stability are considerable. In fact, one could describe the molecular mechanism involved as the protein-equivalent of a Rube Goldberg machine.

A key component of this Rube Goldberg machine is H59. Previously, Sjöberg and co-workers showed that a H59A variant was not able to discriminate effectively between dATP and ATP and implicated H59 in triggering allosteric regulation (Rofougaran et al., 2008). Studies on mammalian RNR, and the equivalent residue (D57), have additionally showed the importance of this residue in allosteric regulation (Caras and Martin, 1988; Reichard et al., 2000). Now, through this work, we can explain how H59 communicates the presence of dATP or ATP and triggers the appropriate response in the *E. coli* class Ia RNR enzyme. We can now also describe that response, which we find involves three sets of conformational changes: the H59-triggered forming/unforming of allosteric effector site 2; the upward/downward tilting of the β-hairpin and accompanying relaxing/tugging of helix 2; and the winding/unwinding of helix 2 and accompanying sealing/revealing of the β subunit binding pocket on α.

In terms of stabilizing the α_4_β_4_ inactive ring-like state, H59 contributes by communicating the presence of dATP to Phe87 and Trp28 such that they adopt positions that block allosteric effector site 2 (**Fig. 10a, Fig. S5**). With allosteric effector site 2 closed for nucleotide binding, dATP in site 1 anchors the tip of the β-hairpin down by contacts made to R10 and K9. With the tip down, a strained helix 2 is unwound, and the binding surface for the β subunit is stabilized (**Fig. 10a,c**).

**Figure 10.**
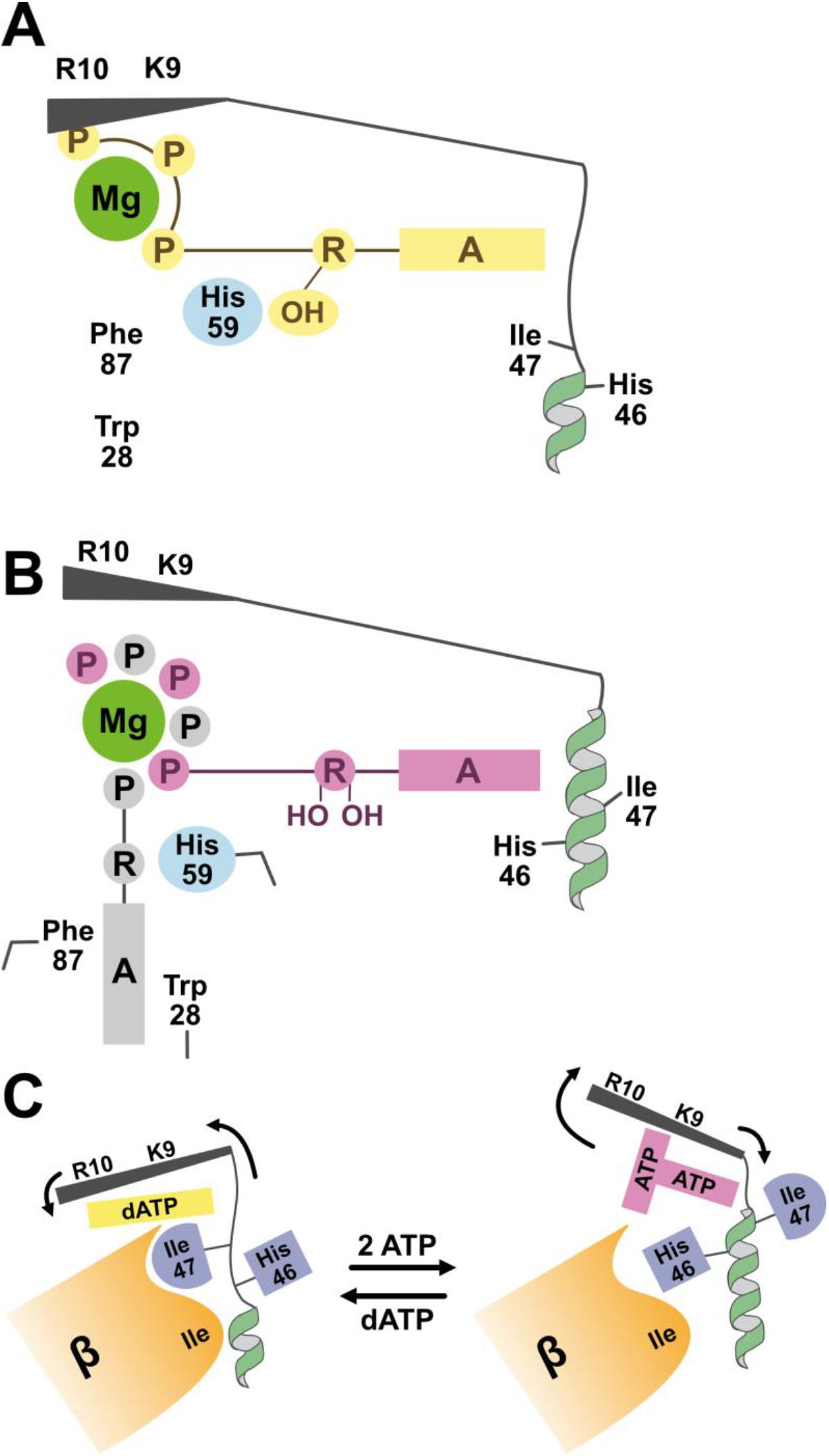
Cartoon representation for the different states of the cone domain. (A) dATP-bound cone domain: H59 hydrogen bonds to the ribose (R) hydroxyl group of dATP (yellow); site 2 is blocked by Phe87 and Trp28; β-hairpin (black) is anchored down via contacts between dATP and R10 and K9; and helix 2 (green) is unwound, creating binding surface for β. (B) ATP-bound cone domain: H59 does not engage in hydrogen bonding to the site 1 ribose (R) of ATP (pink) and H59 is tilted; F87 and W28 have moved and site 2 is open with ATP (grey) bound; the β-hairpin (black) is pushed up due to presence of 2ATPs; the strain on helix 2 (green) is decreased and helix 2 is rewound. (C) The binding surface for β (orange) is alternatively created by dATP (yellow) pulling the β-hairpin (black) down, and hidden by 2 molecules of ATP pushing the β-hairpin (black) up, which unwinds and rewinds helix 2 (green) respectively.

The molecular mechanism by which ATP frees the β subunit from the α_4_β_4_ ring is surprisingly elaborate. With RNR turned off, the ratio of ATP to dATP will increase, and ATP, despite its lower affinity for allosteric effector site 1 (K_d_ of ∼120 μM), will displace dATP (K_d_ of ∼ 6 μM) (Brown and Reichard, 1969b; Ormö and Sjöberg, 1990). Due to the extra hydroxyl group on the ribose of ATP, the ribose sits higher in allosteric effector site 1 to prevent a steric clash between the 2’ hydroxyl group and I22 (see **Fig. 3F**). This repositioning of the ribose, which has also been reported for human RNR (Fairman et al., 2011), breaks the hydrogen bond between H59 and the 3’ hydroxyl group. The side chain of H59 tilts to one side.

The tilting of the H59 side chain starts a Rube Goldberg-like mechanism in which H59 movement results in the movement of the side chain of F87, which in turn results in the movement of the side chain of W28. With the side chains of F87 and W28 repositioned, the second ATP can bind to previously unavailable site 2 (**Fig. 10b**). With six ATP phosphates now positioned around one Mg^2+^ ion, the β-hairpin tip is pushed up, and with the β-hairpin lever tipped up, the strain on helix 2 residues is decreased and helix 2 re-winds, sealing away I47 and the binding pocket for the β subunit (**Fig. 10c**). Ensuring the departure of β, helix 2 re-winding swaps hydrophobic residue I47 with a polar and potentially charged H46. Thus, the Rube Goldberg mechanism concludes with an exchangeable protein surface that alternatively attracts (I47-outward facing) and repels (H46-outward facing) the β subunit. Consistent with the ability of an Ile-to-His exchange at a small interface (∼525 Å^2^) to disrupt that interface, is the previous observation (Chen et al., 2018) that site-directed single substitutions of residues at this interface (e.g. L43Q, S39F, E42K) abolish ring formation.

It is quite impressive from a protein design prospective that a 95-residue domain has two “hideaway” binding sites, one for a nucleotide effector (site 2) and the other for a protein subunit, and that these sites are in communication. The exposure of one binding site necessitates that the other is sealed. When ATP is bound and site 2 is exposed, the β-subunit binding site is tucked away. Conversely, when dATP is bound and site 2 is tucked away, the β-subunit binding site is exposed. Importantly, we were able to demonstrate that site 2 is relevant to allosteric activity regulation through a W28A substitution. Although dATP inhibits W28A RNR in the absence of any ribonucleoside triphosphate, addition of ATP or GTP restores activity for W28A RNR under conditions that are inhibitory for the WT protein. In other words, when site 2 is open, the identity of the nucleotide in site 1 does not matter. Thus, like the H59A RNR variant (Rofougaran et al., 2008), W28A RNR does not discriminate between dATP and ATP. Although unexpected, the involvement of two ATP binding sites in the cone domain of *E. coli* class Ia RNR has a certain structural and chemical logic to it; when ATP levels rise in response to RNR inactivity, that the binding of two ATP molecules per α subunit, rather than one ATP molecule, are required to shift the conformation equilibrium from α_4_β_4_ to α_2_β_2_.

It is too early to say whether human class Ia RNR will utilize one or two ATP molecules in its allosteric regulatory mechanism. W28 is not conserved in human RNR and there is no evidence for the unwinding of helix 2 in the formation of the α−α interface of the dATP-inhibited α_6_ ring (Brignole et al., 2018). D57 (equivalent of H59) appears to be responsible for signaling the presence of dATP versus ATP (Brignole et al., 2018; Caras and Martin, 1988; Reichard et al., 2000), but the molecular response that follows is unknown. Given that the interface involved in formation of inactive rings is different in human RNR (α−α) than in *E. coli* (α−β), we do not expect the molecular mechanism to be the same. This difference in the nature of the inhibited states is exciting and has potential application for selective RNR inhibitor design. FDA-approved drug hydroxyurea and prodrugs gemcitabine and clofarabine either inhibit human RNR and not *E. coli* or both human and *E. coli* class Ia RNR. There are no FDA approved inhibitors that are specific for bacterial RNRs.

Molecules that stabilize the inactive ring structures of RNR would be expected to be successful RNR inhibitors. Notably, clofarabine triphosphate, which is used in the treatment of pediatric acute leukemia (Pession et al., 2010), inhibits human RNR with the generation of “persistent hexamers” (Aye et al., 2012) and does not inhibit the α_4_β_4_-forming *E. coli* class Ia RNR. Recently, we showed that the class Ia RNR from *N. gonorrhea* (*Ng*RNR) forms α_4_β_4_ inactive rings that are analogous to the rings formed by the *E. coli* enzyme, and that compounds that inhibit *Ng*RNR are not cross-reactive with human RNR (Chen et al., 2018). Although there is no structure of *Ng*RNR bound to these compounds, we do know that *N. gonorrhea* develops resistance to them when mutations occur in *Ng*RNR at the α−β interface of the α_4_β_4_ ring, suggesting that these compounds target an inactive ring structure (Chen et al., 2018).

The work presented here should aid in the development of compounds that target, stabilize and thereby increase the lifetime of inactive α_4_β_4_ ring structures of bacterial RNRs. In particular, our studies suggest that small molecules that prevent site 2 from opening, by blocking W28 from moving, for example, would stabilize the inactive α_4_β_4_ ring structure. The inactive ring structure should also be stabilized through maintenance of the β-hairpin in the downward position or maintenance of the unwound conformation of helix 2. In contrast, compounds that stabilize an open site 2 would be expected to impede down-regulation by dATP and lead to persistently active bacterial RNRs. We hope that the molecular information presented here will facilitate the development of new antibiotic compounds. Antibiotic resistance is an eminent threat (CDC, 2018; Willyard, 2017) and RNR is a largely unexplored target to address this threat.

## Materials and Methods

### Reagent preparation

Sodium salts of CDP, ATP, dATP, GTP, and TTP (Sigma-Aldrich) were dissolved into assay buffer (50 mM HEPES pH 7.6, 15 mM MgCl_2_, 1 mM EDTA). The pH was slowly adjusted to 7-8 with NaOH using pH indicator paper and the concentration was determined spectroscopically using ε_271_ of 9.1 mM^-1^ cm^-1^ for CDP, ε_259_ of 15.4 mM^-1^ cm^-1^ for ATP, ε_259_ of 15.2 mM^-1^ cm^-1^ for dATP, ε_253_ of 13.7 mM^-1^ cm^-1^ for GTP, and ε_262_ of 9.6 mM^-1^ cm^-1^ for TTP. For the structures of wild-type α_2_ bound to dATP and ATP, high-purity 100 mM solutions of ATP and dATP were purchased from USB Corporation or Invitrogen. A high-purity 100 mM solution of dGTP was purchased from USB Corporation for ultrafiltration assays. [2,8-^3^H]-ATP tetraammonium salt and [methyl-^3^H]-TTP tetraammonium salt of high activity were obtained from Moravek Biochemicals and added to freshly prepared, unlabeled ATP or TTP, lyophilized, and resuspended in 50 mM HEPES pH 7.6, to give a final activity of 690 or 1070 cpm nmol^-1^ for ^3^H-ATP and 470 cpm nmol^-1^ for ^3^H-TTP. Pall Nanosep 30K Omega filters were used for ultrafiltration assays. All crystallization reagents were purchased from Hampton Research unless otherwise specified.

### Construct and protein preparation

Untagged α_2_ and β_2_ were prepared as previously described (Salowe et al., 1987; Salowe and Stubbe, 1986). The concentrations of α_2_ and β_2_ were determined using ε_280_ of 189 and 131 mM^−1^ cm^−1^, respectively; unless noted otherwise, all molar concentrations correspond to the subunit dimer.

His6-W28A-, F87A-, and F97A-α_2_ were generated from a previously described His_6_-α_2_ construct (**Fig. S9**) in pET28 (Minnihan et al., 2011) by Quikchange mutagenesis (Stratagene) with primers from Integrated DNA Technologies (**Table S5**). His_6_-α_2_ variants were purified as described previously for wild-type His_6_-α_2_ (Minnihan et al., 2011). The His_6_-tag was not removed. For His_6_-W28A-α_2_, an estimated ε_280_ of 182 mM^-1^cm^-1^ was used to determine the final protein concentration.

The α_2_-β_C35_ fusion construct was made by sequential megaprimer mutagenesis (Xu et al., 2003) in which double-headed PCR primers were used to amplify a 144 bp insert encoding the C-terminal tail of β (the C-terminal 35 residues) from a plasmid encoding wild-type β. These megaprimers were subsequently used to amplify the wild-type His_6_-α-encoding plasmid, while excising the final eight residues of the α C-terminal tail. The final construct contains an N-terminal His_6_ tag and thrombin cleavage site, α residues 1-753 (of 761 native residues), and β residues 342-376 (**Fig. S9**). This α_2_-β_C35_ fusion construct was purified as described previously for wild-type His_6_-α_2_ (Minnihan et al., 2011). As the resulting construct has no additional tryptophan or tyrosine residues, ε_280_ of 189 mM^-1^ cm^-1^ was used to determine the final protein concentration.

### Crystallization of wild-type α_2_ with ATP and dATP

Wild-type α_2_-dATP was crystallized by the hanging drop vapor diffusion method. Initial conditions were identified in 96-well sparse matrix screening trays (Qiagen and Hampton Research) dispensed by an Art Robbins Instruments Phoenix liquid-handling robot. For preparation of optimized crystals, 116 μM untagged α_2_ in assay buffer (50 mM HEPES pH 7.6, 15 mM MgCl_2_, 1 mM EDTA), which was also containing 5 mM DTT, and 10 mM dATP was mixed in a 1:1 ratio (drop size 2 μL) with a precipitant solution containing 6.8 % (w/vol) PEG 3350, 80 mM HEPES pH 7.3, 280 mM MgCl_2_, 4 % (vol/vol) glycerol, and 1.0 % cyclohexyl-methyl-β-D-maltoside (CYMAL-1) detergent and equilibrated over a reservoir of 500 μL of precipitant solution at 4 °C. After one week of growth, crystals were cryo-protected by looping through a solution of 10.5 % (w/vol) PEG 3350, 100 mM HEPES pH 7.3, 380 mM MgCl_2_, 13 % (vol/vol) glycerol and plunged directly in liquid N_2_. α_2_-ATP was crystallized as described for α_2_-dATP with the following modifications: the protein concentration was lowered to 88 μM, the PEG 3350 concentration was raised to 8.0 % (w/vol), and all steps were performed at room temperature (∼20°C). After two days of growth, crystals were cryo-protected by looping through a solution of 8.5 % (w/vol) PEG 3350, 100 mM HEPES pH 7.3, 300 mM MgCl_2_, 5 mM DTT, 5 mM ATP and successive concentrations of 10, 15, and 20 % (vol/vol) glycerol and plunged into liquid N_2_.

### Crystallization of W28A-α_2_

Crystals of W28A-α_2_ were identified in sparse matrix trays as described for wild-type α_2_ above. For screening and optimization of the ATP/CDP co-crystal complex, His_6_-W28A-α_2_ at 60 µM in assay buffer (50 mM HEPES pH 7.6, 15 mM MgCl_2_, 1 mM EDTA) was pre-incubated with 10 mM ATP and 1 mM CDP for 20 min at ∼25 °C before mixing with precipitant solutions. Hanging drop optimization trays were set up with a 2 μL drop size in a 1:1 ratio of protein to precipitant at 18 °C. The optimized conditions contained 1.9 M (NH_4_)_2_SO_4_, 4 % (w/vol), PEG MME 500, and 0.1 M bis-Tris pH 6.5. Crystals grew over the course of several weeks and were harvested after approximately one month. Crystals were briefly soaked in a cryo-protectant solution containing 2.5 M (NH_4_)_2_SO_4_, 4 % (w/vol), PEG MME 500, and 0.1 M bis-Tris pH 6.5, 5 mM ATP, 1 mM CDP, and 15 % (vol/vol) glycerol and plunged in liquid nitrogen.

Crystals of W28A-α_2_ for structural analysis with dATP/GTP and dATP/ATP were grown as described above for wild-type α_2_ with minor modifications. After optimization the modified conditions contained 6.5-7.0 % (w/vol) PEG 3350, 0.1 M HEPES 7.0, 0.35 M MgSO_4_, 5 % (vol/vol) glycerol. The His_6_-W28A-α_2_ protein was prepared at 110 µM in assay buffer containing 3-5 mM ATP or 1 mM dATP or both 3-5 mM ATP with 1 mM dATP. Two μL sitting drops with a 1:1 ratio of protein to well solution were prepared at 4 °C and crystals grew over the course of several days and were harvested after 1-2 weeks. Harvested crystals were additionally soaked in a solution containing nucleotides, 1 mM dATP and 3 mM ATP or 1 mM dATP and 3 mM GTP, and cryoprotectant, 10 % (w/vol) PEG 3350, 20 % (vol/vol) glycerol, 0.25 M MgCl_2_, and 0.1 M HEPES pH 7.6, for ∼15 min and then plunged in liquid nitrogen.

### Crystallization of α_2_-β_C35_ with dATP

Crystals of the α_2_-β_C35_ fusion construct co-crystallized with dATP were identified in sparse matrix screens as described above. For screening and optimization, His_6_-α_2_-β_C35_ at 75 µM in assay buffer was pre-incubated with 1 mM dATP and for 20 min at ∼4 °C before mixing with precipitant solutions. Sitting drop optimization trays were prepared with a 2 μL drop size in a 1:1 ratio of protein to precipitant at 4 °C. Optimized conditions contained 1.4 M NaCl and 8 % (w/vol), PEG 6000. Crystals grew over the course of two months and were harvested after approximately three months. Crystals were soaked in a cryo-protectant solution containing 1.4 M NaCl, 15 % (w/vol), PEG 6000, 15 % (vol/vol) glycerol, 15 mM MgCl_2_, 50 mM HEPES pH 7.6, and 1 mM EDTA with 1 mM dATP and 1 mM CDP for approximately 2 min and then plunged in liquid nitrogen. No CDP was observed bound in this crystal form, consistent with the observation that a large shift of the barrel is required for substrate binding (Zimanyi et al., 2016).

### X-ray data collection

Diffraction data for wild-type α_2_-dATP and α_2_-ATP were collected at the Advanced Photon Source (APS) on beamline 24ID-C on a Quantum 315 CCD detector at 100 K. The W28A-α_2_-(ATP)_2_/CDP, W28A-α_2_-(dATP/ATP), W28A-α_2_-(dATP/GTP), and wild-type α_2_-β_C35_-dATP datasets were collected at APS beamline 24ID-C on a Pilatus 6M detector (Dectris) at 100 K. Diffraction data were indexed, integrated, and scaled using HKL2000 (Otwinowski and Minor, 1997), with statistics shown in **Table S1**.

### Structure solution and refinement

The method of structure solution is described individually for each structure below. Model refinement statistics for wild-type α_2_ structures are found in **Table S2** and model refinement statistics for W28A α_2_ structures are found in **Table S3**. For all structures, multiple rounds of refinement were performed with phenix.refine (Adams et al., 2010) in the SBGRID software package (Morin et al., 2013). For all structures, refinement consisted of rigid body, positional, and individual B factor refinement. Translation-libration-screw (TLS) B factor refinement was used for all structures except W28A-α_2_-(ATP)_2_/CDP. Manual rebuilding and geometry correction was performed in Coot (Emsley et al., 2010). Simulated annealing composite omit maps calculated in Phenix were used to validate modeling of ligands in all structures. For structures with resolution of 3 Å or better, waters were placed automatically in Phenix with manual editing and placement in Coot (Emsley et al., 2010). Ligand restraint files were obtained from the Grade Web Server (Global Phasing Ltd.). Coordination distances for Mg^2+^ ions were explicitly defined at 2.1 Å with loose restraints. All structural figures were made in PyMOL v. 1.7 and v. 2.3.7 (Schrödinger, LLC). Refinement statistics for each final model are given in **Table S2** (wild-type cone domain) and **Table S3** (W28A mutants).

The α_2_-dATP co-crystal structure was solved by molecular replacement in Phaser (McCoy et al., 2007) at 2.55-Å resolution using a single α monomer (chain A) from a previously solved structure of α_2_ (PDB ID 3R1R) (Eriksson et al., 1997) as the search model. The α_2_-ATP co-crystal structure was solved by molecular replacement in Phaser at 2.62-Å resolution using the refined α_2_-dATP co-crystal structure with all ligands and waters removed. Since the crystal form of these two structures is essentially identical, the cross-validation sets for the α_2_-ATP structure was preserved from the α_2_-dATP structure. For both structures, the resolution was extended to the full range after partial model building and refinement at lower resolution. CNS 1.3 (Brünger et al., 1998) was used for early stages of refinement. ATP or dATP and Mg^2+^ ions were placed into omit density prior to addition of water molecules. In both α_2_-dATP and α_2_-ATP models, two α chains are present in the asymmetric unit in a physiological dimer. Loose non-crystallographic symmetry (NCS) restraints on both coordinate positions and B factors were used throughout refinement and then removed during the final rounds. Residues 5-736 (of 761) are present in each chain of both structures. 1-2 additional residues are visible in some chains at the N-terminus, but are poorly structured. The C-terminal 25 residues are thought to form a flexible tail essential for the re-reduction of the active site disulfide upon turnover.

The W28A-α_2_-(ATP)_2_/CDP structure was solved by molecular replacement in the Phenix implementation of Phaser (McCoy et al., 2007) with a 3.60-Å resolution cutoff. Due to the large changes in the overall conformation upon binding of the substrate CDP, the best search model was a single α_2_ dimer from the wild-type α_4_β_4_-dATP/CDP structure (PDB ID 5CNS) (Zimanyi et al., 2016). The initial R_free_ for this molecular replacement solution was 0.41. The resolution was extended to 2.60 Å after rigid body refinement and manual rebuilding of the model. Simulated annealing and real-space refinement were used early on until the model converged. Eight α_2_ dimers are present in the asymmetric unit, organized as two separate dimer-of-dimer units. A stable α_4_ oligomeric state has never been observed in solution for *E. coli* class Ia RNR, and analysis of the overall structure by the PISA server suggests that the only stable assembly is the α_2_ dimer and not α_4_. The overall structure of the α subunit’s (β/α)_10_ barrel closely resembles that observed in the substrate/effector-bound α_4_β_4_ complex, despite the complete absence of β_2_ from this crystal form. The cone domain was partially deleted and rebuilt manually during refinement. ATP and Mg^2+^ ions were placed into omit density. NCS restraints were used throughout refinement. Composite omit maps were used to verify the final structure, especially to ensure model bias did not influence the rebuilding of the cone domain. The N-terminal His_6_ tag and thrombin cleavage site and residues 737-760 at the C-terminus are disordered in all chains. Residues 645-652, which form a flexible β-hairpin, are poorly ordered in four of the eight chains and have been omitted where there is no clear density.

The W28A-α_2_-dATP/ATP and W28A-α_2_-dATP/GTP structures were solved by molecular replacement in Phaser (McCoy et al., 2007) with the wild-type α_2_-(ATP)_2_ structure as the model with no resolution cutoff. This crystal form contains lattice contacts that are similar to those observed in the wild-type α_2_-ATP and α_2_-dATP structures, but four molecules are present in the asymmetric unit instead of two, and the length of the b-axis is increased by 20 Å (a ∼17% increase). The C-terminal tail of α (24 residues) is disordered in all of the chains of each structure along with 3-4 residues at the N-terminus, and the N-terminal His_6_ tag and thrombin cleavage site. There is no clear reason for the change in space group that is apparent from the crystal packing, but it is possible that the presence of the N-terminal His_6_ tag disrupts some crystal contacts in this crystal form, although the N-terminus is not ordered in either W28A or wild-type free α_2_ structures. To prevent overfitting, strict NCS for both coordinate positions and B factors was maintained throughout refinement. All ligands were removed prior to molecular replacement and were rebuilt based on omit maps. Due to the low resolution and similarity of the starting model, few structural changes were observed during refinement. dATP was placed in the specificity site based on omit maps. ATP/dATP and GTP/dATP were placed according to the positions of the ATP/ATP pair in the wild-type α_2_-(ATP)_2_ structure. To assess whether the structure was phase-biased by the choice of α_2_-(ATP)_2_ as the molecular replacement model, α_2_-dATP was also tested. The molecular replacement solution was substantially worse in this case, but could be corrected with rigid body refinement of the hairpin and rotamer flips of W28 and F87. Composite omit maps were used to verify ligand placement, the conformation of F87, and the W28A mutation. The N-terminal His_6_ tag and thrombin cleavage site and residues 737-760 at the C-terminus are disordered in all chains. All other residues are present in all four chains of both structures.

The α_2_-β_C35_-dATP/CDP structure was solved to 2.10-Å resolution by molecular replacement in Phaser (McCoy et al., 2007) using an α_2_ structure that contained two bound peptides that mimicked the sequence of the β tail (PDB ID 1R1R) (Eriksson et al., 1997) with no resolution cutoff. The first 20 residues of α_2_ were not included in the search model. The initial R_free_ for this starting model was 0.29. The crystal form contains two molecules in the asymmetric unit. Only one of the two cone domains is fully structured (beginning after the N-terminal His_6_ tag and thrombin cleavage site); the other cone domain has no density for residues 1-19 and 47-58 due to a close crystal contact. Residues 738-753 of the α tail and 342-364 (β numbering) of the attached β tail are disordered, but the remainder of the β tail, 365-376 (β numbering), renumbered as 1365-1376, is bound as previously observed for both the β peptide in the α_2_ structure and for the β tail in the α_4_β_4_ complex (Eriksson et al., 1997; Zimanyi et al., 2016). No NCS restraints were used as there were substantial differences in the cone domains in the two protomers. dATP was modeled into the final structure based on omit maps. At both specificity and activity sites, a Mg_2+_ ion coordinates three phosphate oxygens of dATP and three water molecules. Composite omit maps generated in Phenix (Adams et al., 2010) were used to verify the cone domain conformations and effector ligands.

### Determination of equilibrium ATP binding parameters by ultrafiltration

The ultrafiltration method described by Ormö and Sjöberg (Ormö and Sjöberg, 1990) was used with modifications to determine the equilibrium binding parameters for ATP. In place of Millipore Ultrafree-MC filter units with polysulfone PTTK membranes, which are no longer available, we used Pall Nanosep 30 kDa molecular weight cutoff filters with polyethersulfone membranes. For experiments with ^3^H- ATP, filters were washed twice by adding 200 μL equilibration buffer (50 mM HEPES pH 7.6, 15 mM MgCl_2_, 1 mM EDTA, 5 mM DTT, 0.5 mM cold ATP) to the filter unit, equilibrating in a 25 °C water bath for 5 min, then spinning in a table top centrifuge at 12,000 ×g for 30 s. After the second wash, the sample consisting of 7-20 μM α_2_ (in assay buffer with 5 mM DTT) and 50-1000 μM ^3^H- ATP with a specific activity of either 690 cpm nmol^-1^ or 3090 cpm nmol^-1^ in a total volume of 150 μL was added to the filter. The solution was equilibrated in a 25 °C water bath for 5 min. and a 25 μL aliquot was taken for determination of total nucleotide concentration. The sample was then centrifuged at 12,000 × g for 1 min. 25 μL of the filtrate was then taken to determine free nucleotide concentration by scintillation counting. To isolate the binding events at the activity site, 100 or 500 μM dGTP was included in the sample before equilibration. The amount of bound nucleotide was found by subtracting the amount of free nucleotide from the total nucleotide. Data were plotted as a saturation binding curve and analyzed using non-linear regression and a one- site specific binding model in the program Prism (GraphPad).

### Determination of α_2_ activity

The activity of wild-type α_2_ and the W28A, F87A, and F97A α_2_ variants was determined in a spectrophotometric coupled assay as previously described (Ge et al., 2003). *E. coli* thioredoxin (TR) and thioredoxin reductase (TRR) were prepared as previously described (Chivers et al., 1997; Zimanyi et al., 2016). All reaction mixtures contained 30 µM TR, 0.5 µM TRR, 200 µM NADPH, and 1 mM CDP as substrate in assay buffer (defined above). Standard active and inactive conditions contained either 3 mM ATP or 175 µM dATP as allosteric effectors, respectively. Concentrations of nucleotides used in titration experiments are given in the **Fig. 8** legend. Linear fitting of the initial rates was performed in the Cary WinUV Kinetics program (Varian/Agilent) and data were plotted in MATLAB (Mathworks).

## Acknowledgements

We thank Yasmin Chau, and Victor Cruz for assistance in identifying and optimizing crystallization conditions, and Yimon Aye for assistance with analysis of the ultrafiltration data. This work was supported by National Institutes of Health (NIH) Grant R35 GM126982 (C.L.D), T32GM08334 (C.M.Z.) and P30-ES002109, and a National Science Foundation Graduate Research Fellowship grant no. 0645960 (M.A.F.). C.L.D. is a HHMI Investigator. Portions of this work were performed at the Advanced Light Source (ALS) at the Berkeley Center for Structural Biology beamline, which is supported in part by the National Institutes of Health (NIH), National Institute of General Medical Sciences, and the Howard Hughes Medical Institute (HHMI). ALS is supported by the U.S. Department of Energy (DOE) under Contract No. DE-AC02-05CH11231. In addition, this work is based upon research conducted at the Northeastern Collaborative Access Team beamlines, which are funded by the NIH (P30 GM124165). The Pilatus 6M detector on 24-ID-C beam line is funded by a NIH-ORIP HEI grant (S10 RR029205). The Eiger 16M detector on 24-ID-E is funded by a NIH-ORIP HEI grant (S10OD021527). This research used resources of the Advanced Photon Source, a U.S. Department of Energy (DOE) Office of Science User Facility operated for the DOE Office of Science by Argonne National Laboratory under Contract No. DE-AC02-06CH11357.

---

**Figure S1.**
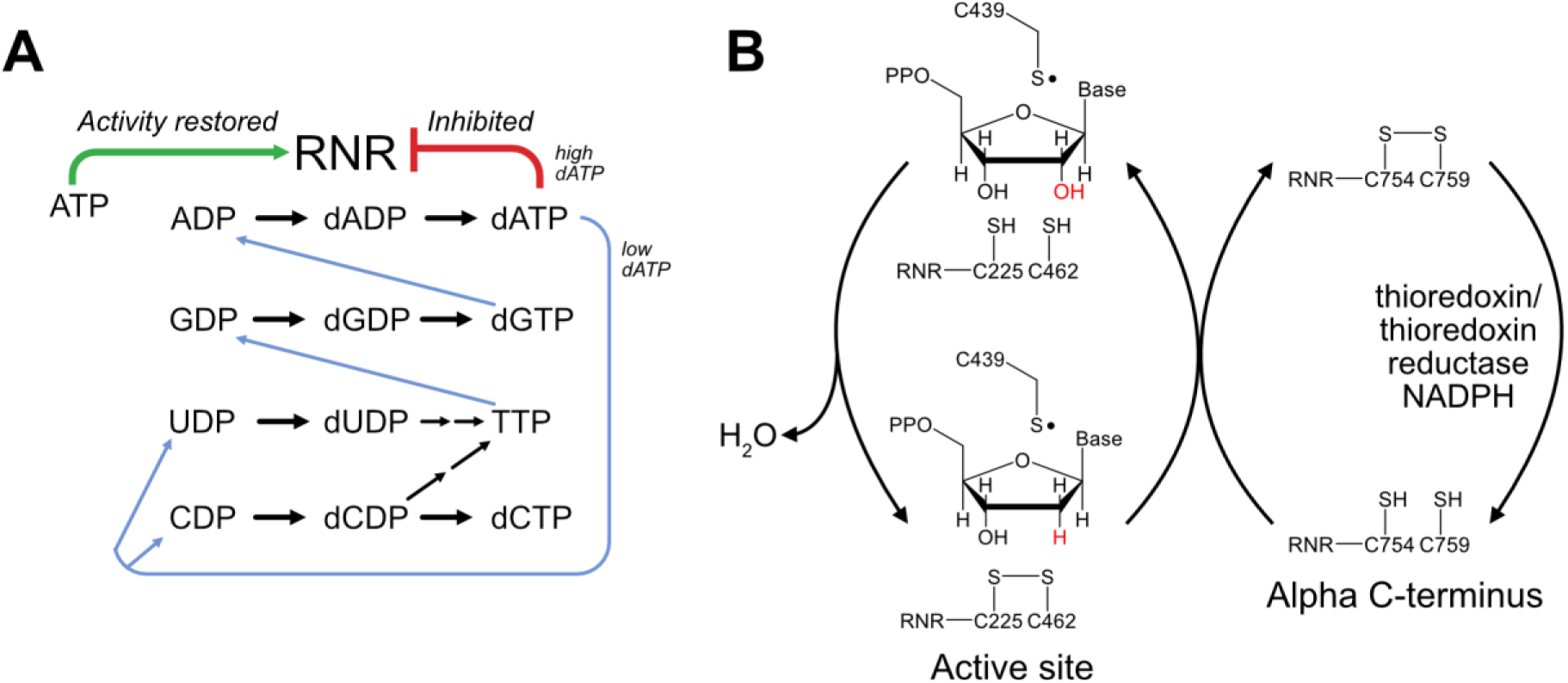
Class Ia RNRs reduce all four ribonucleotide substrates and are allosterically regulated. (A) Substrate specificity is regulated allosterically via the binding of deoxynucleotides. Under low concentrations of dATP, CDP and UDP reduction is favored. TTP in turn leads to GDP reduction and dGTP leads to ADP reduction. When the concentration of dATP increases, it can bind to a second allosteric site, the activity site, which inhibits enzymatic activity. ATP can compete for binding at this activity site to restore activity. (B) Enzymatic turnover in class Ia RNR involves a series of redox-active cysteine pairs. The reducing equivalents for nucleotide reduction are initially provided by a pair of cysteines (C225/462) in the active site that is oxidized to form a disulfide concomitant with product formation. This disulfide is reduced by a second pair of redox-active cysteines (C754/759) found at the C-terminus of the α_2_ subunit. Ultimately, the disulfide between C754 and C759 is reduced via the thioredoxin/thioredoxin reductase pair together with NADPH, thus allowing for additional rounds of turnover.

**Figure S2.**
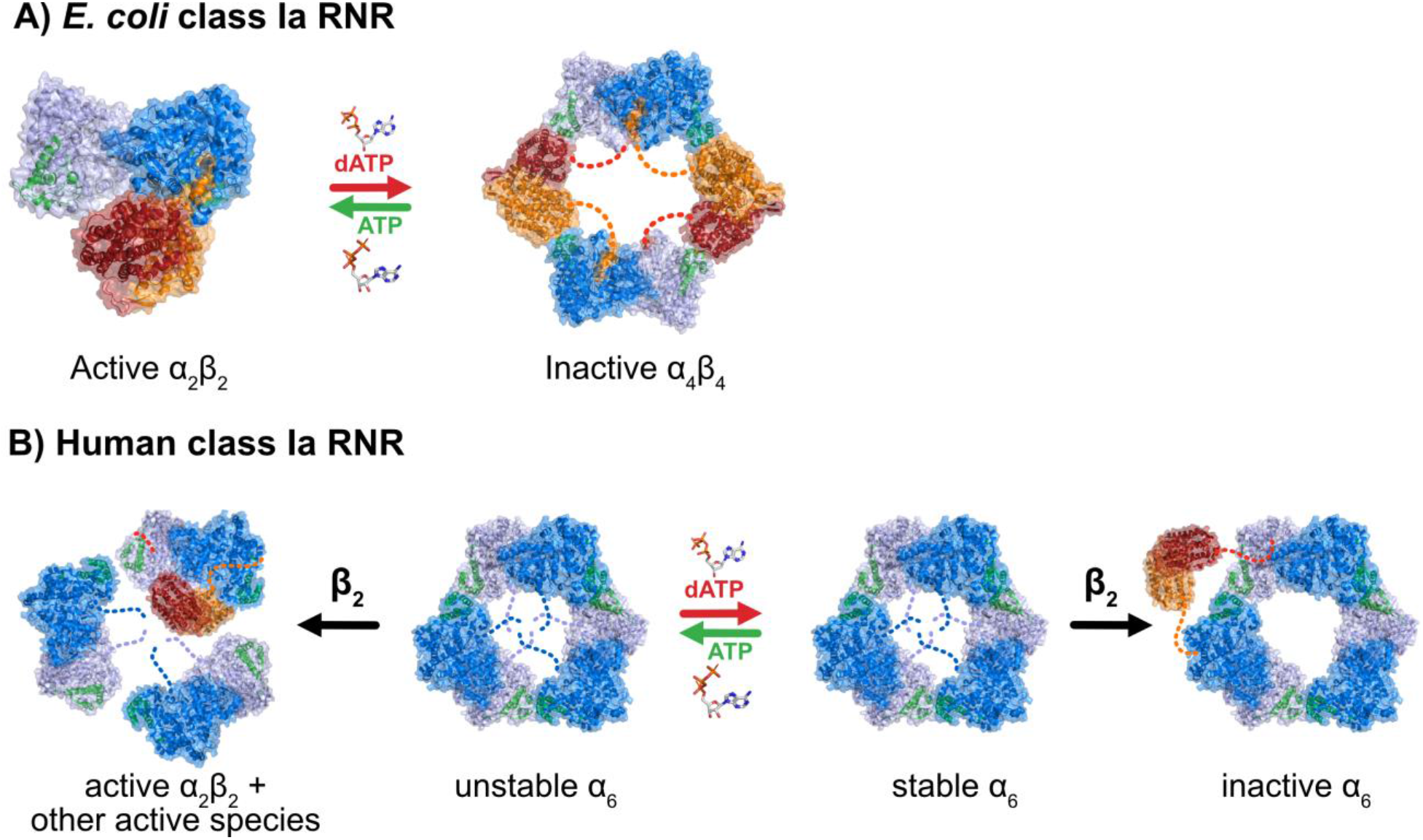
Oligomeric changes in *E. coli* and Human class Ia RNR. (A) *E. coli* class Ia RNR transitions from an active complex (PDB: 6W4X) in the presence of ATP to an inactive α_4_β_4_ ring (PDB: 5CNS) in the presence of dATP. (B) In the presence of ATP, human class Ia RNR forms an unstable α_6_ complex, along with other species involving the β_2_ complex. In the presence of dATP, human class Ia RNR forms an inactive, stable α_6_ complex that is unable to position the β_2_ complex in the correct catalytic position (α_6_ PDB: 6AUI, β_2_ PDB: 2UW2).

**Fig. S3.**
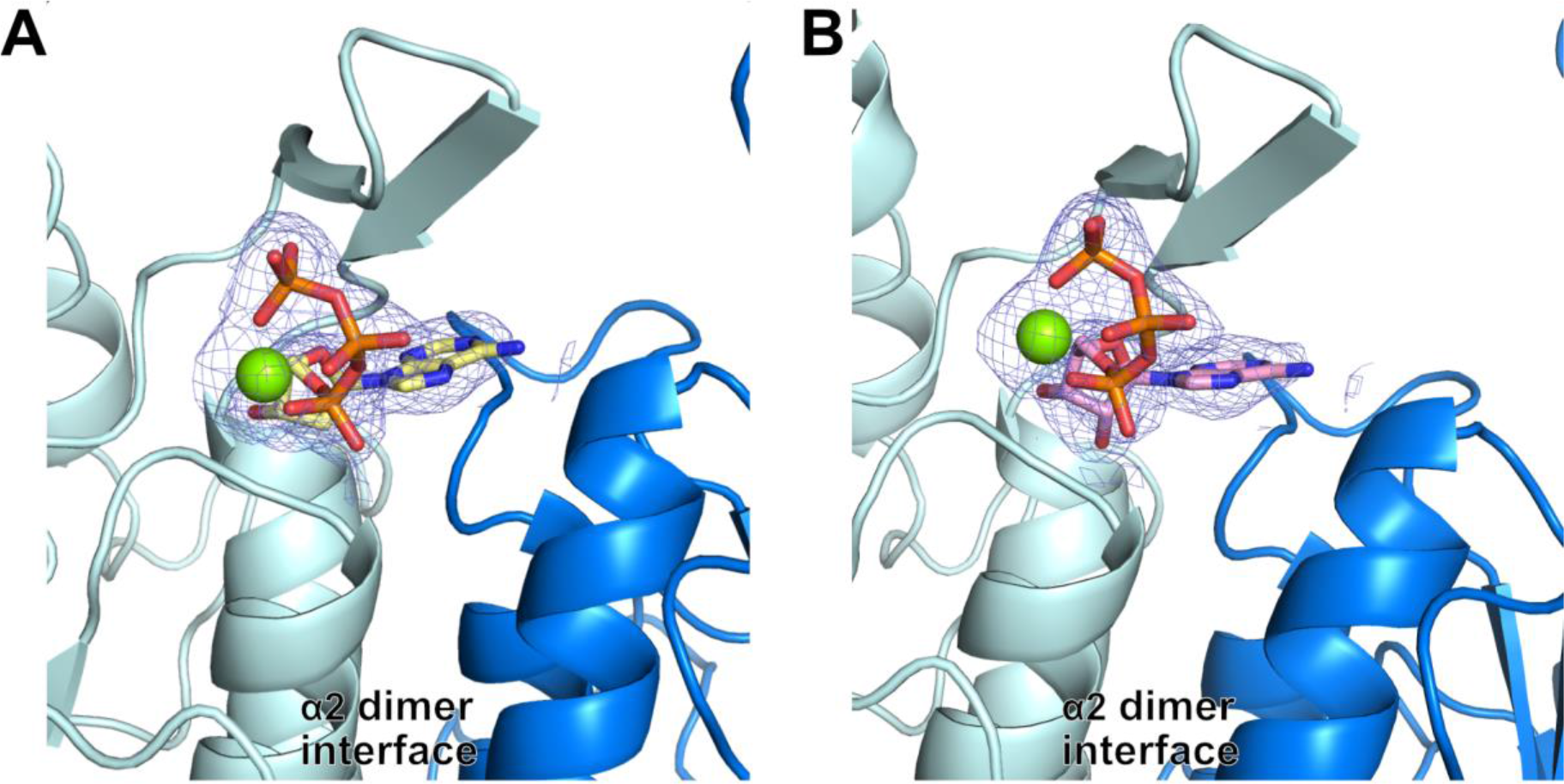
Omit electron density for ATP and dATP binding in the allosteric specificity site of α_2_ structures of *E. coli* class Ia RNR. (A) dATP (yellow) bound at one of the specificity sites in WT α_2_ dimer that has dATP also bound in the activity site. One monomer is shown in a darker shade of blue. (B) ATP (pink) bound at one of the specificity sites in WT α_2_ dimer that has 2 ATPs also bound in the activity site. One monomer is shown in a darker shade of blue. Composite omit density contoured at 1.0 σ is shown in blue mesh.

**Figure S4.**
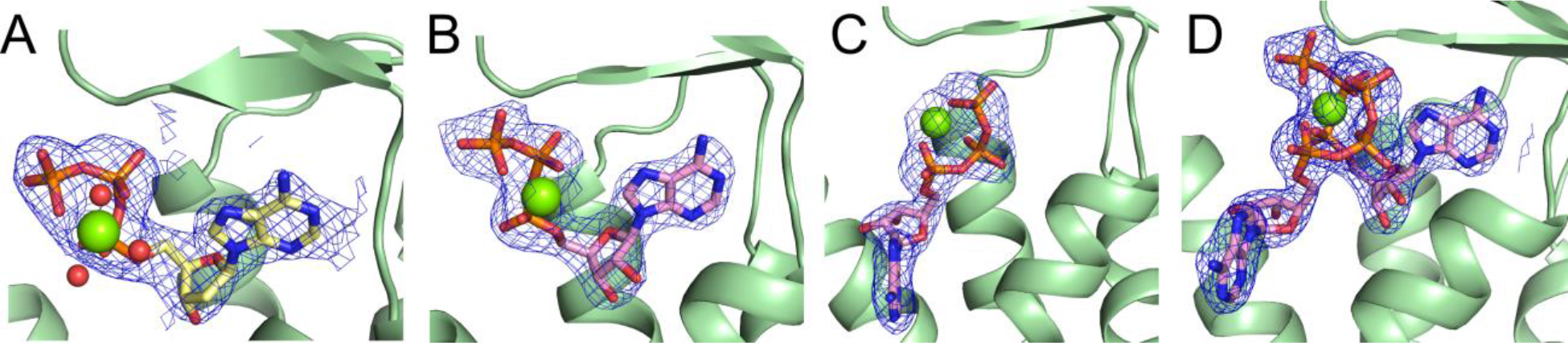
Omit electron density for ATP and dATP binding in the allosteric activity site in the cone domain of α_2_ structures of *E. coli* class Ia RNR site. (A) dATP in site 1 of WT α_2_-dATP. (B) ATP in site 1 of WT α_2_-(ATP)_2_. (C) ATP in site 2 of WT α_2_-(ATP)_2_. (D) Both ATP molecules in the cone domain of WT α_2_-(ATP)_2_. Composite omit density contoured at 1.0 σ is shown in blue mesh. Site 1 and site 2 are explained in the text.

**Figure S5.**
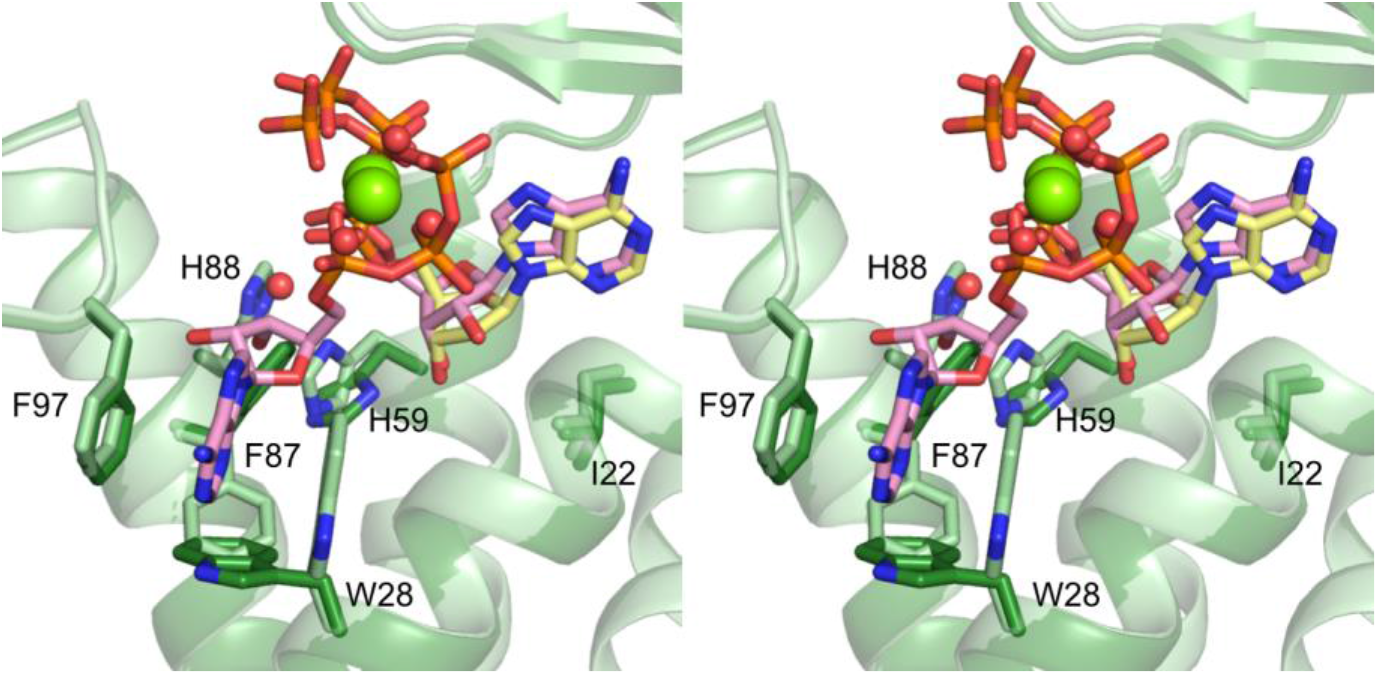
Stereoview of dATP-α_2_ structure superimposed with ATP-α_2_ structure. Stereoview of α_2_- dATP (dark green, dATP molecule in yellow) aligned with α_2_-(ATP)_2_ (light green, ATP molecules in pink).

**Figure S6.**
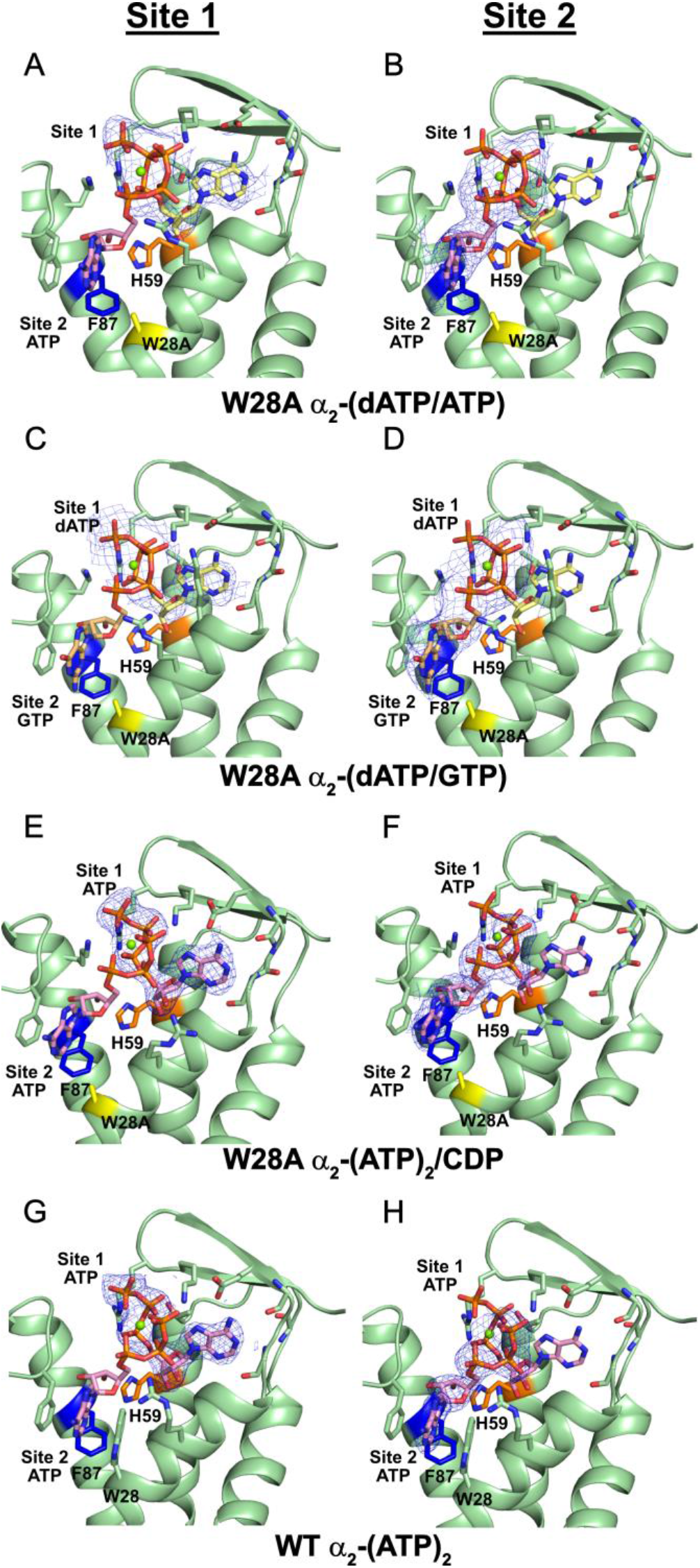
Omit electron density for activity effector binding in W28A structures compared to WT. (A) dATP bound in site 1 of W28A α_2_-(dATP/ATP). (B) ATP bound in site 2 of W28A α_2_-(dATP/ATP). (C) dATP (yellow carbons) bound in site 1 of W28A α_2_-(dATP/GTP). (D) GTP (tan carbons) bound in site 2 of W28A α_2_-(dATP/GTP). (E) ATP (pink carbons) bound in site 1 of W28A α_2_-(ATP)_2_/CDP. (F) ATP bound in site 2 of W28A α_2_-(ATP)_2_/CDP. (G) ATP bound in site 1 of WT α_2_-(ATP)_2_. (H) ATP bound in site 2 of WT α_2_-(ATP)_2_. All site 1 residues, including residues F87 highlighted in blue and H59 highlighted in orange, in the W28A constructs are identical to WT. Composite omit density contoured at 1.0 σ is shown in blue mesh, and the W28A mutation is highlighted in yellow. Nucleotides bound in the other site not focused on in a panel are shown in transparent sticks.

**Fig. S7.**
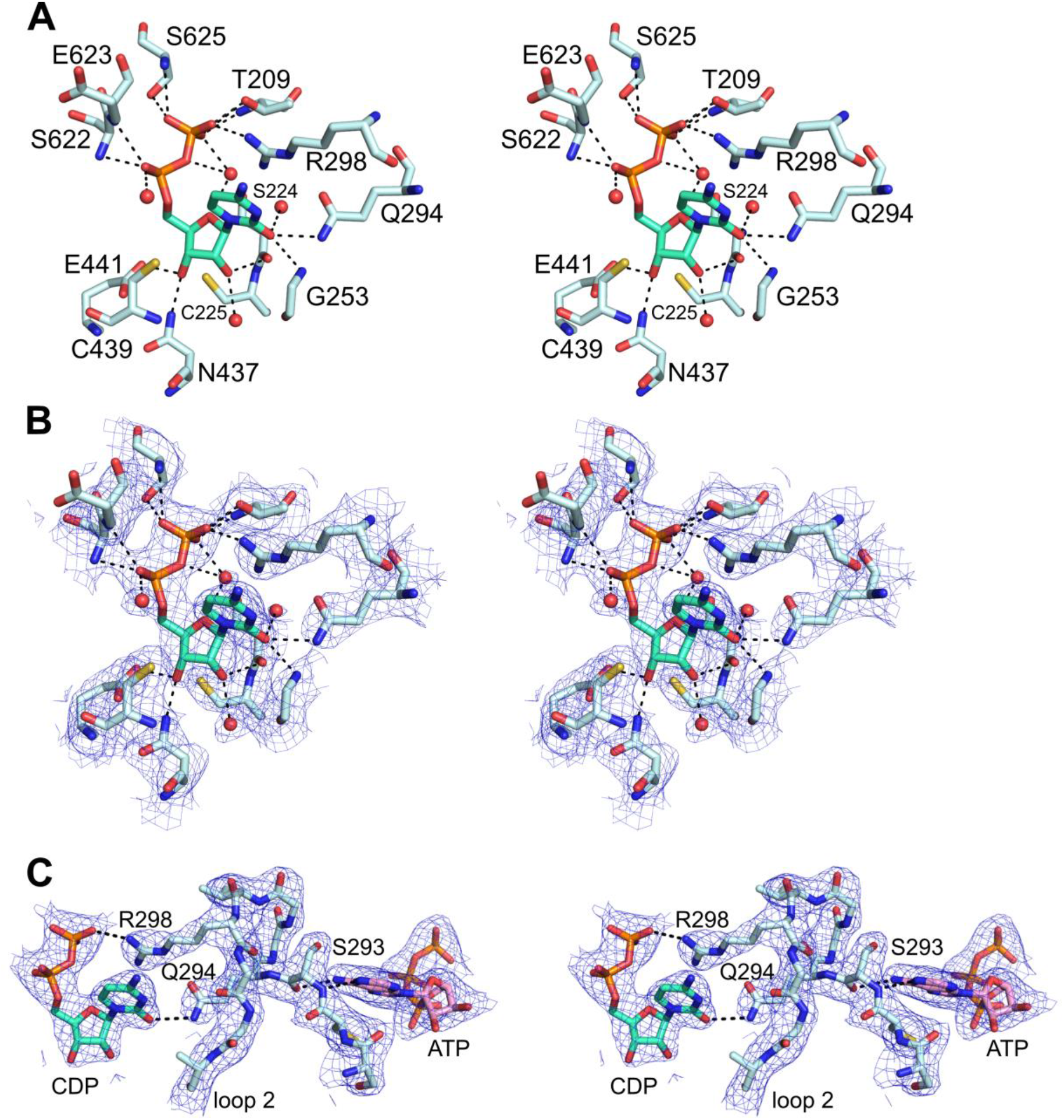
Stereoviews of CDP bound in the active site of W28A α_2_-(ATP)_2_ /CDP. (A) Wall-eyed stereoview of CDP (cyan) interaction with protein in the active site of W28A α_2_-(ATP)_2_/CDP. Hydrogen bonding interactions shown with black dashed lines. (B) Wall-eyed stereoview of composite omit electron density for CDP structure shown in A. Omit density contoured at 1.0 σ in blue mesh. (C) Wall-eyed stereoview of CDP’s (cyan) interaction with the specificity loop (labelled loop 2) and specificity effector ATP’s (pink) interaction with loop 2. As was observed previously (Zimanyi et al., 2016), hydrogen bonds (black dash lines) between ATP and Ser293 backbone atoms of loop 2 orient the side chain of Q294 into the active site to hydrogen bond with the base of CDP. R298 from loop 2 contacts the CDP phosphates.

**Figure S8.**
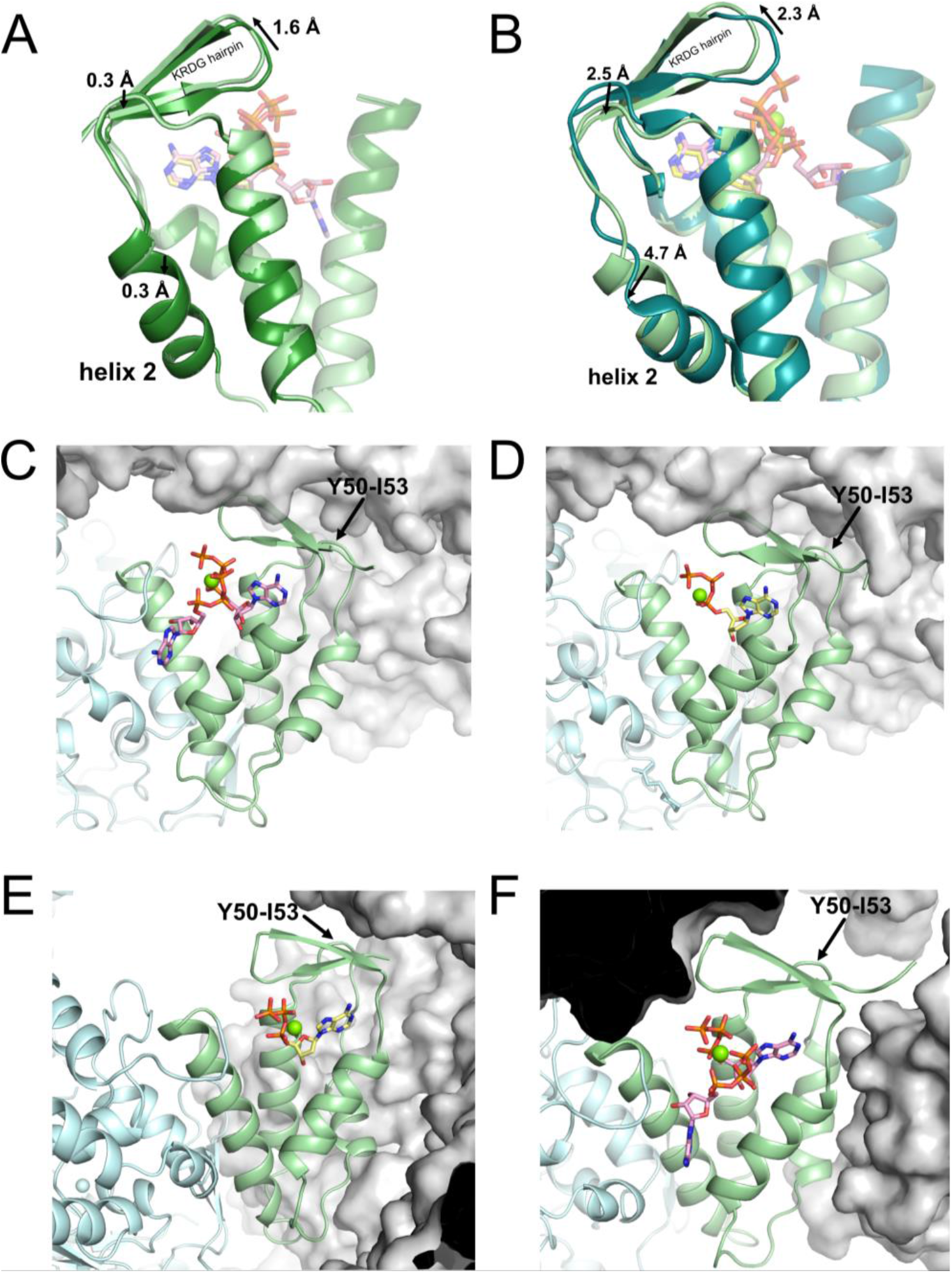
Lattice contacts affect β-hairpin movement and thus the unwinding of helix 2 of the cone domain. (A) Virtually no helix 2 unwinding is observed when α_2_-(ATP)_2_ (light green) and α_2_-(dATP) (dark green) are compared. (B) Helix 2 unwinding seen in alignment of α_2_-(ATP)_2_ (light green) and α_4_β_4_-(dATP) (PDB: 5CNS, dark teal). This panel is the same as Fig. 9B. (C-F) Lattice contacts of α_2_ structures shown with symmetry mates displayed in grey surfaces. Cone domain is colored light green, rest of protein is colored light blue, dATP is shown in yellow and ATP is shown in pink. Residues Y50-I53 that are at the base on the β-hairpin are highlighted with black arrow. (C) α_2_-(ATP)_2_, (D) α_2_-(dATP), (E) α_2_-βC35-(dATP) (F) W28A α_2_-(ATP)_2_

**Figure S9.**
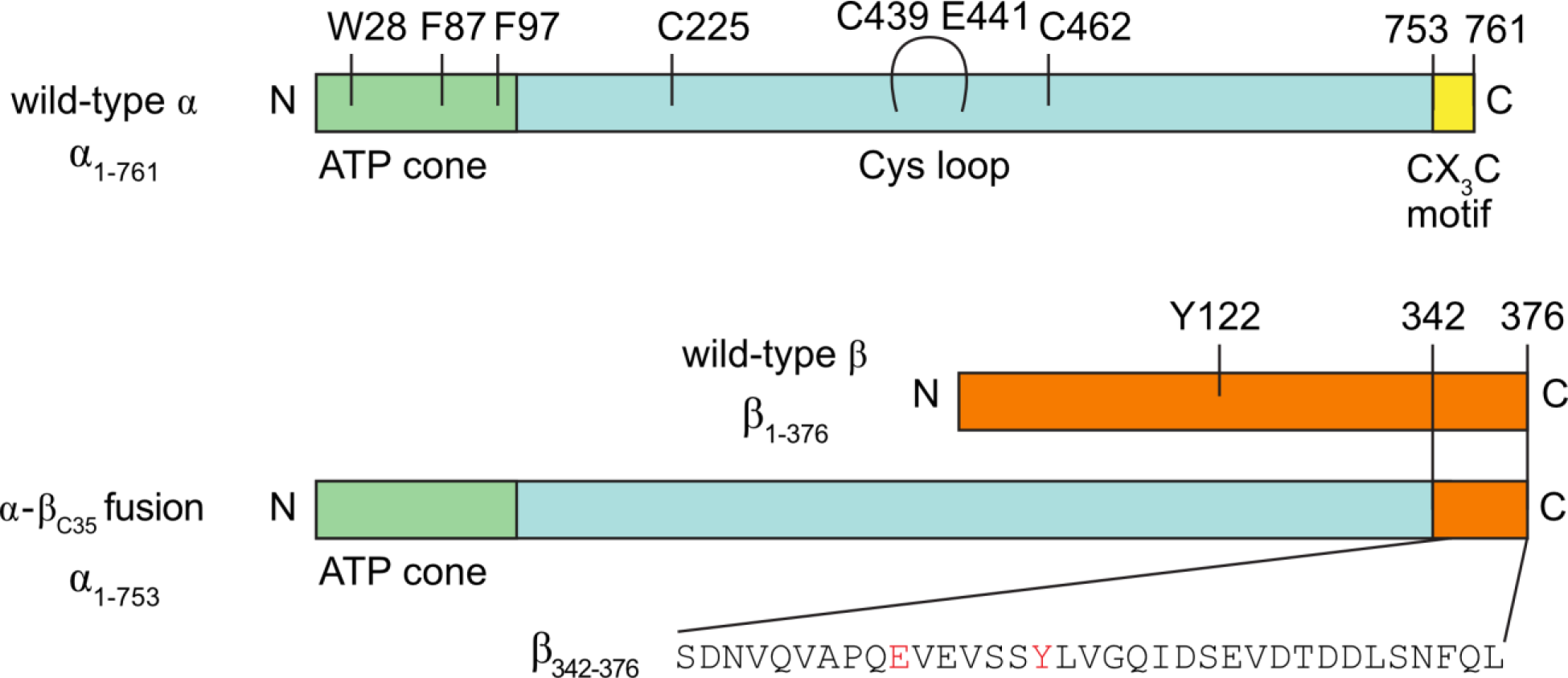
Schematic diagram of constructs used in this study. Residues chosen for substitution via site-directed mutagenesis within the cone domain are labeled as well as residues known to be involved in radical transfer or catalysis.

**Table S1.** X-ray data collection statistics.

| | $\alpha_2$-dATP | $\alpha_2$-(ATP)<sub>2</sub> | $\alpha_2$-$\beta_{C35}$-dATP |
| --- | --- | --- | --- |
| Space group | P4<sub>3</sub>2<sub>1</sub>2 | P4<sub>3</sub>2<sub>1</sub>2 | P2<sub>1</sub>2<sub>1</sub>2<sub>1</sub> |
| Cell dimensions (a, b, c; Å) | 118.5, 118.5, 290.3 | 118.9, 118.9, 290.9 | 88.2 111.2 209.7 |
| Wavelength (Å) | 0.9793 | 0.9794 | 0.9792 |
| Resolution (Å) | 50.0-2.55<br>(2.59-2.55) | 50.0-2.62<br>(2.67-2.62) | 50-2.10<br>(2.14-2.10) |
| R<sub>sym</sub><sup>a</sup> | 0.083 (0.830) | 0.107 (0.772) | 0.083 (0.801) |
| <I/σI><sup>a</sup> | 13.7 (2.0) | 12.9 (2.7) | 15.4 (2.7) |
| Completeness (%)<sup>a</sup> | 99.9 (100.0) | 99.4 (99.5) | 98.7 (99.0) |
| Unique reflections | 68422 | 63413 | 118973 |
| Redundancy | 4.8 (4.9) | 7.2 (7.1) | 8.9 (7.9) |
| CC1/2 | 0.694 | 0.756 | 0.79 |

| | W28A $\alpha_2$-(ATP)<sub>2</sub>/CDP | W28A $\alpha_2$-(dATP/ATP) | W28A $\alpha_2$-(dATP/GTP) |
| --- | --- | --- | --- |
| Space group | C222<sub>1</sub> | P2<sub>1</sub>2<sub>1</sub>2<sub>1</sub> | P2<sub>1</sub>2<sub>1</sub>2<sub>1</sub> |
| Cell dimensions (a, b, c; Å) | 288.7 316.6 158.9 | 112.7 139.9 300.8 | 113.5 141.0 301.8 |
| Wavelength (Å) | 0.9792 | 0.9792 | 0.9792 |
| Resolution (Å) | 50.0-2.60<br>(2.64-2.60) | 50.0-3.40<br>(3.52-3.40) | 50.0-3.55<br>(3.68-3.55) |
| R<sub>sym</sub><sup>a</sup> | 0.128 (>1) | 0.094 (0.792) | 0.075 (0.938) |
| <I/σI><sup>a</sup> | 12.7 (1.2) | 9.8 (1.8) | 11.9 (1.6) |
| Completeness (%)<sup>a</sup> | 99.8 (99.9) | 98.8 (97.1) | 95.9 (93.6) |
| Unique reflections | 218946 | 65872 | 57221 |
| Redundancy | 7.2 (6.4) | 3.8 (3.6) | 4.2 (4.0) |
| CC1/2 | 0.54 | 0.54 | 0.65 |

<sup>a</sup>Values for data in the highest resolution bin are shown in ().

**Table S2.** Model refinement statistics for wild-type α_2_ and α_2_-β_C35_ structures.

| | $\alpha_2$-dATP | $\alpha_2$-(ATP)<sub>2</sub> | $\alpha_2$-$\beta_{C35}$-dATP |
| --- | --- | --- | --- |
| Resolution (Å) | 50.0-2.55<br>(2.58-2.55) | 50.0-2.62<br>(2.66-2.62) | 50.0-2.10<br>(2.12-2.10) |
| Number of reflections |  |  | 118834 |
| R<sub>work</sub> | 0.165 | 0.166 | 0.154 |
| R<sub>free</sub> | 0.205 | 0.209 | 0.188 |
| R<sub>free</sub> set size (%) | 3376 (4.9 %) | 3103 (4.9 %) | 3712 (3.1 %) |
| Model components |  |  |  |
| Protein chains | 2 | 2 | 2 |
| Mg<sup>2+</sup> molecules | 4 | 4 | 3 |
| dATP molecules | 4 | -- | 3 |
| ATP molecules | -- | 6 | -- |
| Other ligand molecules | 4<sup>a</sup> | 4<sup>a</sup> | 9<sup>b</sup> |
| Water molecules | 676 | 758 | 1357 |
| Average B factors (Å<sup>2</sup>) |  |  |  |
| Protein | 51.2 | 47.1 | 47.6 |
| Mg<sup>2+</sup> molecules | 62.6 | 45.6 | 41.9 |
| dATP (activity site 1) | 72.3 | -- | 54.6 |
| dATP (specificity site) | 43.2 | -- | 36.0 |
| ATP (activity site 1) | -- | 62.4 | -- |
| ATP (activity site 2) | -- | 79.1 | -- |
| ATP (specificity site) | -- | 48.8 | -- |
| Other ligands | 99.2<sup>a</sup> | 91.4<sup>a</sup> | 58.5<sup>b</sup> |
| Water | 57.9 | 51.0 | 55.6 |
| Coordinate error (Å) | 0.27 | 0.27 | 0.21 |
| R.M.S deviations |  |  |  |
| Bond length (Å) | 0.002 | 0.007 | 0.003 |
| Bond angle (°) | 0.660 | 0.853 | 0.729 |
| Rotamer outliers (%) | 0.16 | 0.16 | 0.48 |
| Ramachandran plot (%) |  |  |  |
| Favored | 97.88 | 97.74 | 97.80 |
| Additionally allowed | 1.85 | 2.12 | 2.06 |
| Disallowed | 0.27 | 0.14 | 0.14 |

<sup>a</sup> two molecules of MES and two molecules of glycerol<sup>b</sup> two molecules of glycerol, six chloride ions, and one sodium ion

**Table S3.** Model refinement statistics for W28A α_2_ structures.

| | W28A $\alpha_2$-(ATP)<sub>2</sub>/<br>CDP | W28A $\alpha_2$-<br>(dATP/ATP) | W28A $\alpha_2$-<br>(dATP/GTP) |
| --- | --- | --- | --- |
| Resolution (Å) | 50-2.60<br>(2.67-2.60) | 50-3.40<br>(3.45-3.40) | 50-3.55<br>(3.59-3.59) |
| Number of reflections | 218879 | 65402 | 56607 |
| $R_{\text{work}}$ | 0.189 | 0.200 | 0.208 |
| $R_{\text{free}}$ | 0.210 | 0.228 | 0.245 |
| $R_{\text{free}}$ set size | 2000 (0.9 %) | 3272 (5.0 %) | 5641 (10.0 %) |
| Model components |  |  |  |
| Protein chains | 8 | 4 | 4 |
| Mg<sup>2+</sup> molecules | 16 | 8 | 8 |
| dATP molecules | -- | 8 | 8 |
| ATP molecules | 24 | 8 | -- |
| CDP molecules | 8 | -- | -- |
| GTP molecules | -- | -- | 4 |
| Other ligand molecules | 40<sup>b</sup> | 0 | 0 |
| Water molecules | 1224 | 0 | 0 |
| Average B factors (Å<sup>2</sup>) |  |  |  |
| Protein | 54.2 | 125.7 | 145.7 |
| Mg<sup>2+</sup> molecules | 53.3 | 85.9 | 113 |
| dATP (activity site 1) | -- | 114 | 167 |
| dATP (specificity site) | -- | 80.6 | 121 |
| ATP (activity site 1) | 59.8 | -- | -- |
| ATP (activity site 2) | 56.8 | 125 | -- |
| ATP (specificity site) | 47.2 | -- | -- |
| GTP (activity site 2) | -- | -- | 178 |
| CDP (substrate) | 44.8 | -- | -- |
| Other ligands | 58.3<sup>b</sup> | -- | -- |
| Water | 51.1 | -- | -- |
| Coordinate error (Å) | 0.31 | 0.37 | 0.48 |
| R.M.S deviations |  |  |  |
| Bond length (Å) | 0.004 | 0.002 | 0.002 |
| Bond angle (°) | 0.762 | 0.581 | 0.618 |
| Rotamer outliers (%) | 0.36 | 0.04 | 0.44 |
| Ramachandran plot (%) |  |  |  |
| Favored | 97.91 | 96.68 | 97.74 |
| Additionally allowed | 1.95 | 2.74 | 2.02 |
| Disallowed | 0.14 | 0.58 | 0.24 |

<sup>a</sup> four molecules of MES and four molecules of glycerol<sup>b</sup> forty sulfate ions

**Table S4.** Structural similarity between the cone domain of selected *E. coli* class Ia RNR crystal forms. All atom RMSD for residues 6-99 is given in angstroms. Only a single α chain was used for each structure.

| | nucleotide free $\alpha_2$<br>(1R1R) | a4b4-dADP/dATP/CDP<br>(5CNS) | $\alpha_2$-(ATP)<sub>2</sub> | $\alpha_2$-dATP | $\alpha_2$-$\beta_{C35}$-dATP | W28A $\alpha_2$-dATP | W28A $\alpha_2$-(ATP)<sub>2</sub> | W28A $\alpha_2$-(GTP/dATP) | W28A $\alpha_2$-(ATP/dATP) |
| --- | --- | --- | --- | --- | --- | --- | --- | --- | --- |
| nucleotide free $\alpha_2$ | | 1.8 | 0.8 | 0.9 | 0.9 | 1.7 | 1.3 | 0.8 | 0.8 |
| $\alpha_4\beta_4$-dADP/dATP/CDP | 1.8 | | 1.8 | 1.7 | 1.7 | 0.8 | 1.0 | 1.8 | 1.8 |
| $\alpha_2$-(ATP)<sub>2</sub> | 0.8 | 1.8 | | 0.2 | 0.5 | 1.7 | 1.1 | 0.2 | 0.2 |
| $\alpha_2$-dATP | 0.9 | 1.7 | 0.2 | | 0.5 | 1.7 | 1.1 | 0.3 | 0.3 |
| $\alpha_2$-$\beta_{C35}$-dATP | 0.9 | 1.7 | 0.5 | 0.5 | | 1.7 | 1.0 | 0.6 | 0.6 |
| W28A $\alpha_2$-dATP | 1.7 | 0.8 | 1.7 | 1.7 | 1.7 | | 1.3 | 1.7 | 1.6 |
| W28A $\alpha_2$-(ATP)<sub>2</sub> | 1.3 | 1.0 | 1.1 | 1.1 | 1.0 | 1.3 | | 1.1 | 1.1 |
| W28A $\alpha_2$-(GTP/dATP) | 0.8 | 1.8 | 0.2 | 0.3 | 0.6 | 1.7 | 1.1 | | 0.1 |
| W28A $\alpha_2$-(ATP/dATP) | 0.8 | 1.8 | 0.2 | 0.3 | 0.6 | 1.6 | 1.1 | 0.1 | |

